# Amyloid pathology compresses dynamic range and degrades spatial coding in an Alzheimer’s mouse model

**DOI:** 10.1101/2025.06.27.661987

**Authors:** Mary Ann Go, Kathrine E. Clarke, Yimei Li, Seigfred V. Prado, Beatriz R. F. Teixeira, Mingyang Gao, Jess J. Yu, Simon R. Schultz

## Abstract

Alzheimer’s disease (AD) disrupts neural circuits vital for memory and cognition. Using two-photon calcium imaging in behaving 5xFAD mice, we examined how amyloid pathology alters hippocampal CA1 activity and spatial coding. We found elevated baseline activity but reduced locomotion-driven firing, leading to a diminished neuronal dynamic range. To our knowledge, this is the first direct experimental evidence for reduced dynamic range in an AD model. These abnormalities were strongest near amyloid plaques and became more widespread with age. We also observed altered network synchrony, degraded spatial coding and increased neuronal response variability. Furthermore, place fields emerged more slowly in both familiar and novel environments, indicating impaired recall and learning. By showing a link between local plaque pathology and impaired flexible modulation of CA1 activity and progressive deficits in spatial memory coding, our study offers new insights into the circuit basis of cognitive decline in AD.

## 1. Introduction

Impairment of memory and cognition is a prominent symptom of Alzheimer’s disease (AD), with spatial and episodic memory processes particularly affected. While substantial advances have been made in characterizing the molecular and neuropathological correlates of AD (Selkoe, 2001), therapeutic progress depends critically on understanding how these pathological signatures lead to alterations in the ability of neural circuits to process, store and recall information.

Disruption of the balance of excitation and inhibition has been shown in AD (Palop et al., 2007; Palop and Mucke, 2016; Van Nifterick et al., 2023; Fortel et al., 2023) and may directly affect circuit function, providing a primary mechanism for cognitive deficits. However, crucial aspects of this puzzle remain poorly understood. For instance, in anesthetized mice, neuronal hyperactivity in hippocampal circuits has been reported (Šišková et al., 2014) especially in the vicinity of amyloid plaques (Busche et al., 2012). However, other studies in behaving mice have reported conflicting results, including little change or even a relative decrease (hypoactivity) in the firing of hippocampal cells (Jun et al., 2020; Mably et al., 2017; Lin et al., 2022; Zhang et al., 2023). Reconciling this discrepancy is key to advancing our understanding of the circuit-level effects of AD. At the same time, it is important to understand how aberrant excitability affects neural circuit function and leads to cognitive deficits.

The hippocampal CA1 circuit provides a good model system to study this problem, as the hippocampus is important for memory function in both humans and rodents (Squire, 2004), it is one of the earliest locations to show amyloid pathology (Hyman et al., 1984), and the CA1 subfield contains many “place cells” - cells tuned for a particular spatial location in an environment (O’Keefe and Dostrovsky, 1971) that remap their spatial tuning in distinct environments (Muller and Kubie, 1987), providing a circuit mechanism for spatial memory.

We leveraged the strengths of two-photon calcium imaging to track the activity of hundreds of neurons per mouse as well as visualize amyloid plaque deposits. Head-fixation is currently required for high quality, stable two-photon imaging in behaving animals (although this may change in the near future; see Zong et al., 2022). Two-photon calcium imaging has been used to map place fields in head-fixed mice navigating a virtual reality (VR) environment (Harvey et al., 2009; Hainmueller and Bartos, 2018), but the use of VR environments using only distal visual cues to study spatial cognition in rodents has been criticized (Aghajan et al., 2015). The limitation of impaired spatial selectivity in head-fixed animals has been largely ameliorated by the development of “floating cage” environments involving multi-modal, proximal cues (Go et al., 2021). *In vivo* two-photon microscopy thus now provides us with the ability to study the how the storage and retrieval of cellular contributions to spatial memory is affected by the three-dimensional distribution of amyloid plaques in the vicinity of individual neurons.

In this study, we examined the relationship between aberrant excitability, the three-dimensional spatial proximity of amyloid plaques, and spatial memory as read out from the hippocampal CA1 place cell network. We used the 5xFAD mouse model, a well-established model of amyloidosis with five human familial AD mutations, and in which high levels of intracellular A*β* begin to accumulate at 1.5 months of age, with plaque deposition beginning around 2 months, and being widespread in the cortex and hippocampus by 6 months (Oakley et al., 2006). We investigated the progression of spatial information coding deficits during maturation and aging of 5xFAD mice, and the extent to which disruptions were synchronous across the circuit, as opposed to affecting cells independently, and how these deficits affected learning of new spatial memories versus the recalling of old.

## 2. Results

To study how amyloid plaques influence the relationship between neural circuit activity and spatial memory, we used a floating real-world environment behavioural apparatus (Go et al., 2021), together with a multi-photon microscope, to image the activity of populations of hippocampal CA1 neurons in head-fixed mice trained to run around a circular track (Fig. 1a) decorated with patterned visuotactile cues. Hippocampal neurons in 5xFAD and wild-type (WT) mice were labeled with GCaMP6s or jGCaMP7s calcium sensors (Fig. 1b). We studied two age groups, the first based on the age at which accumulation of A*β* starts, and the second driven by the age at which plaque deposition becomes widespread (Oakley et al., 2006). Mice in the young group were 2.1 to 3.0 months old at the time of the first imaging session (median 2.7), and mice in the older group were 6.4 to 10.1 months of age (median 7.5). Mice were trained to navigate the circular track for 6-7 days and then imaged. The proximity of amyloid plaques to each neuron was mapped approximately one week after the first neuronal activity imaging session (see Methods for full schedule). Neural activity was extracted using non-negative matrix factorization (Fig. 1c; see Methods) and by deconvolving trains of events using the OASIS algorithm (Friedrich et al., 2017). Activity rates are calculated by accumulating the number of calcium events within a certain time window, and dividing by window duration.

**Figure 1:**
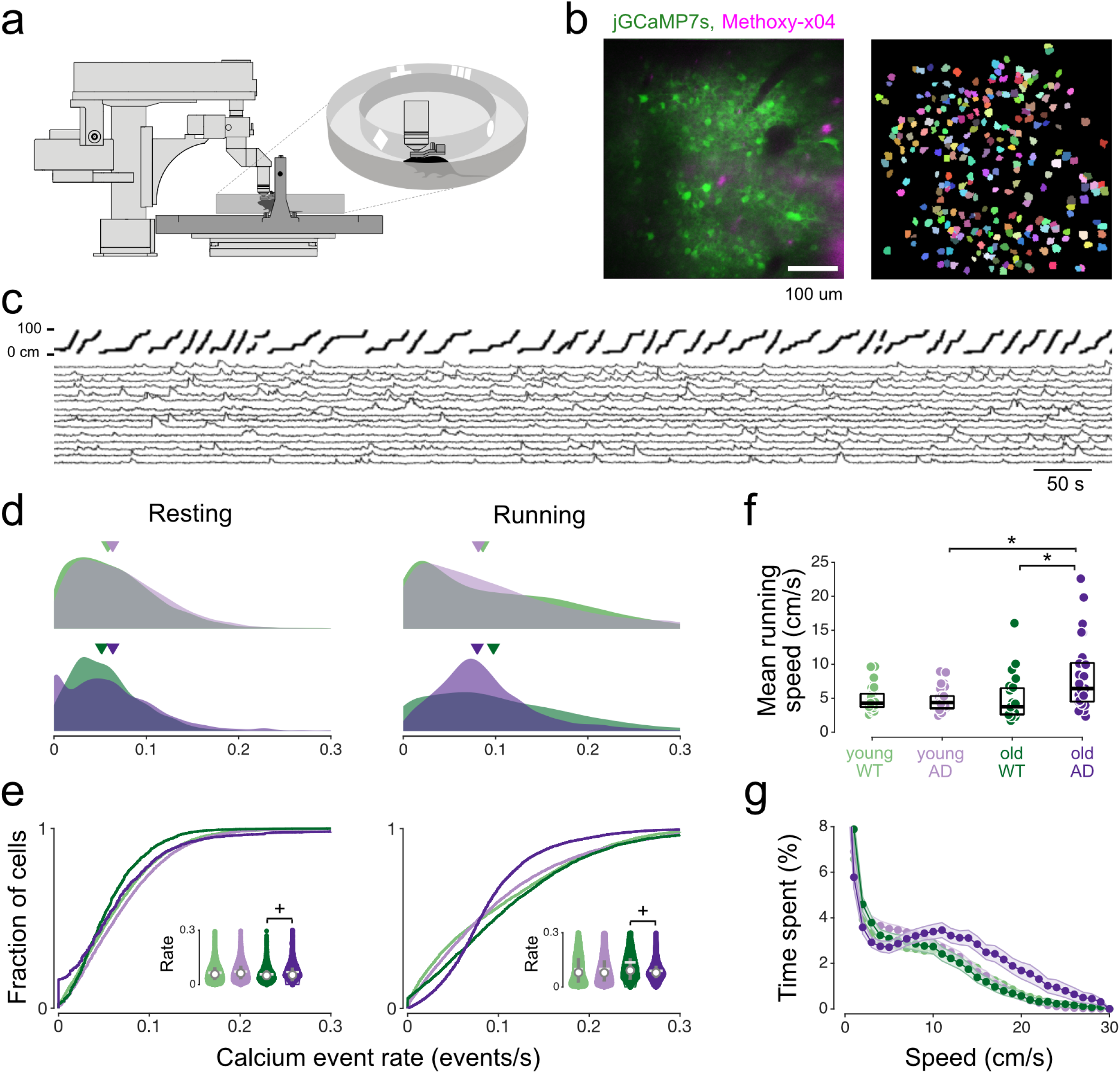
Elevated baseline activity and suppressed firing during locomotion in old 5xFAD mice. **a** Behavioural apparatus: head-fixed mice traverse a floating circular track. **b** (*Left*) Typical FOV showing neurons (green, jGCaMP7s) overlaid with the amyloid plaques (magenta, Methoxy-X04) within 100 vertical microns of the calcium imaging plane. This example from a 7.4-month old 5xFAD mouse. (*Right*) 267 segmented ROIs from image on left. **c** (*Top*) Linearized position of mouse along outer circumference of track; (*Bottom*) Ca^2+^ transients for 15 (of 267) cells from B. See also Fig. S1. **d** Calcium event rate distributions for 5xFAD and WT mice during resting (speed *<* 1.0 cm/s) and running (speed *>* 2 cm/s) periods. Inverted triangles denote median. Group color codes as in (f). Resting, young WT: 0.065 *±* 0.002 events/s (mean *±* SEM); young 5xFAD: 0.072 *±* 0.007 events/s; old WT: 0.056 *±* 0.005 events/s; old 5xFAD: 0.069 *±* 0.006 events/s. Running, young WT: 0.099 *±* 0.009 events/s; young 5xFAD: 0.100 *±* 0.008 events/s; old WT: 0.136 *±* 0.030 events/s; old 5xFAD: 0.090 *±* 0.006 events/s. **e** Cumulative histograms and violin plots (inset) of the distributions in D. Violin plots show mean (white solid line), median (white circle) and interquartile range (gray vertical line). **f** Average running speed during the first 50 laps of each session. Each data point indicates one running session. Boxplot edges denote quartiles while black bars denote median. Young WT: 4.81 *±* 0.47 cm/s; young 5xFAD: 4.84 *±* 0.56 cm/s; old WT: 4.78 *±* 0.85 cm/s; old 5xFAD: 8.14 *±* 1.46 cm/s. **g** Time spent (as a fraction of recording time) at each speed. For **d**-**e**, young WT: n = 5,168 cells pooled from 6 animals, 33 sessions; young 5xFAD: n = 4,925 cells, 6 animals, 27 sessions; old WT: n = 3,475 cells, 8 animals, 27 sessions; old 5xFAD: n = 7,332 cells, 6 animals, 31 sessions. All statistics were performed with hierarchical bootstrap analysis (see Methods). ^+^*p <* 0.050, \**p <* 0.025.

### 2.1 Elevated CA1 baseline firing and suppressed firing during locomotion in old 5xFAD mice

Hippocampal CA1 activity is modulated by both spatial location and running speed (Góis and Tort, 2018). Because activity during rest differs from activity during locomotion — for instance, place cells can spontaneously replay previous trajectories during immobility (Ó lafsdóttir et al., 2018) - we analyzed neuronal activity during stationary and locomotor periods separately.

During rest, defined as periods when speed of movement did not exceed 1.0 cm/s, neural activity was higher in old 5xFAD mice than in old WT mice, as shown in Fig. 1d,e (see also Fig. S2). This baseline hyperactivity was not evident in young mice. It is notable that in old AD mice, the activity distribution showed a mode at zero, suggesting an increased number of nearly silent cells.

We next asked whether this shift in baseline activity was also present during locomotion, when CA1 activity is expected to increase because of both place selectivity and speed modulation. We analysed periods when mice were running (periods in which speed exceeded 2 cm/s). Across all groups, calcium event rates were higher during running than during rest (Fig. 1d,e, right panels). Similar to the resting phase, young 5xFAD mice showed no substantial difference from young WT mice. In contrast, old 5xFAD mice showed significantly lower locomotion-induced activity than old WT mice. Their calcium event rate distribution was also narrower: low event rates shifted upward, whereas very high event rates were less frequent, indicating a diminished range of activity.

Given the strong differences in neuronal activity distributions observed during running, and that motor deficits have previously been observed in 5xFAD mice by 9 months of age (Jawhar et al., 2012), we also examined running speed. We found that old 5xFAD mice ran faster, on average, than old WT mice (Fig. 1f), whereas the young groups had comparable speeds. In addition, old 5xFAD mice spent more time moving at higher speeds (Fig. 1g). Thus, the old 5xFAD mice showed behavioral hyperactivity despite reduced locomotion-related neural activity.

### 2.2 Reduced dynamic range of activity in 5xFAD mice during locomotion

The capability of neurons to discriminate stimulus intensities is given by its dynamic range (Gollo et al., 2012). The combination of elevated baseline activity and reduced locomotion-related firing in old 5xFAD mice suggests a reduction in neuronal dynamic range, with potential consequences for information coding. To examine this further, we calculated speed tuning curves for the average population activity, as well as for individual neurons. The population speed tuning curve in old 5xFAD mice started at a higher baseline but was flatter compared to the other groups (Fig. 2a), indicating reduced dynamic range of the population signal. To quantify this effect at the single neuron level, we defined the dynamic range of firing for each cell as the ratio between the maximum and non-zero minimum values of its speed tuning curve. We refer to this measure as the speed-related dynamic range, noting that neuronal activity increases above baseline during running do not necessarily reflect speed tuning alone, because place cells may also fire when the mouse passes through their place field, or due to other triggers. Speed-related dynamic range in single neurons was significantly lower in old 5xFAD mice than in old WT mice (Fig. 2b), but relatively unchanged between young WT and young 5xFAD mice. Dynamic range was also lower in old 5xFAD mice than in young 5xFAD mice.

**Figure 2:**
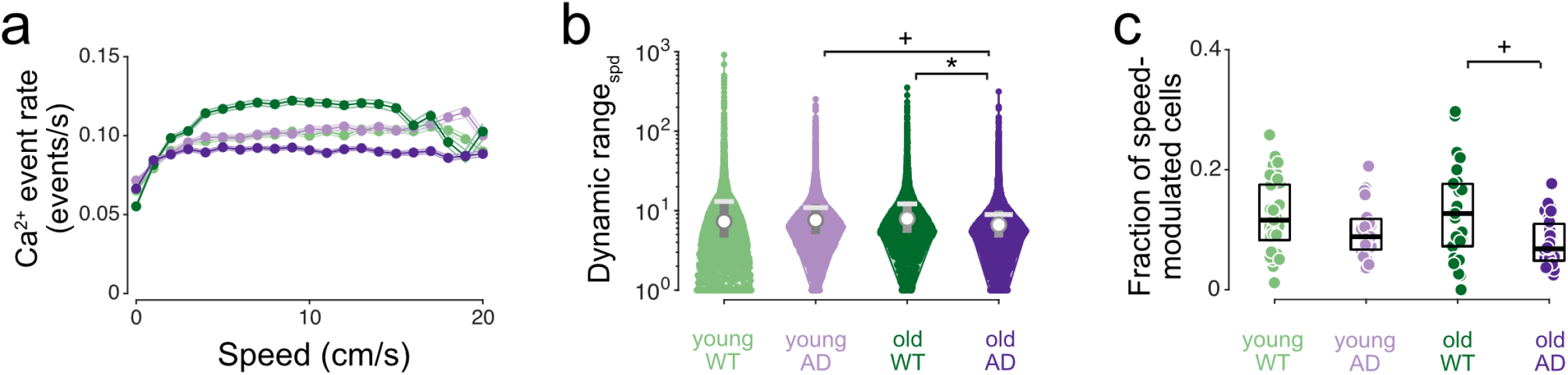
Reduced neuronal dynamic range during locomotion in old 5xFAD mice. **a** Population average speed tuning curves. Group color codes as in (b). **b** Violin plots of dynamic range of neuronal activity during locomotion, defined as the ratio of the maximum and non-zero minimum points on the speed tuning curve for each cell. Young WT: 13.05 *±* 1.78; young 5xFAD: 10.92 *±* 0.85; old WT: 12.18 *±* 1.24; old 5xFAD: 8.93 *±* 0.92. ^+^*p* = 0.048, \**p* = 0.022. **c** Proportion of cells in each group showing significant speed modulation. Each data point indicates a session. Young WT: 0.14 *±* 0.03; young 5xFAD: 0.12 *±* 0.02; old WT: 0.16 *±* 0.03; old 5xFAD: 0.08 *±* 0.01. ^+^*p* = 0.032. See also Figs. S2 and S3. All statistics were performed with hierarchical bootstrap analysis. For all panels, young WT: n = 5,310 cells pooled from 6 animals, 33 sessions; young 5xFAD: n = 4,955 cells, 6 animals, 27 sessions; old WT: n = 3,517 cells, 8 animals, 27 sessions; old 5xFAD: n = 7,400 cells, 6 animals, 31 sessions.

To examine speed tuning further, we asked whether the fraction of cells modulated by speed was affected by age or genotype. Old 5xFAD mice had a significantly smaller fraction of speed-modulated cells than old WT mice (Fig. 2c). We identified three different types of modulation profiles: cells positively modulated by speed, cells negatively modulated by speed, and cells with maximal firing at a preferred speed below 70% of the maximum animal speed (Fig. S3a,b). Across groups, positively modulated cells were the most common profile, whereas negatively modulated cells were the least common (Fig. S3c). The relative proportions of these profiles were broadly similar across groups, although subtype-specific differences were observed in old 5xFAD mice. Old 5xFAD mice had a lower proportion of positively modulated cells and a higher proportion of negatively modulated compared to old WT and young 5xFAD mice. Old 5xFAD mice also had a higher proportion of preferred-speed cells compared to young 5xFAD mice.

### 2.3 Neuronal circuit alterations begin in the vicinity of amyloid plaques

A key advantage of our *in vivo* two-photon microscopy approach is that it allows amyloid plaques to be visualized in three dimensions and related to the properties of nearby neurons. We therefore examined whether the neuronal circuit alterations we observed (such as elevated baseline firing and reduced dynamic range) varied with distance from amyloid plaques (Fig. 3a). We averaged the quantity of interest, such as calcium event rate, across all cells at a certain distance from a plaque, then averaged across all plaques.

**Figure 3:**
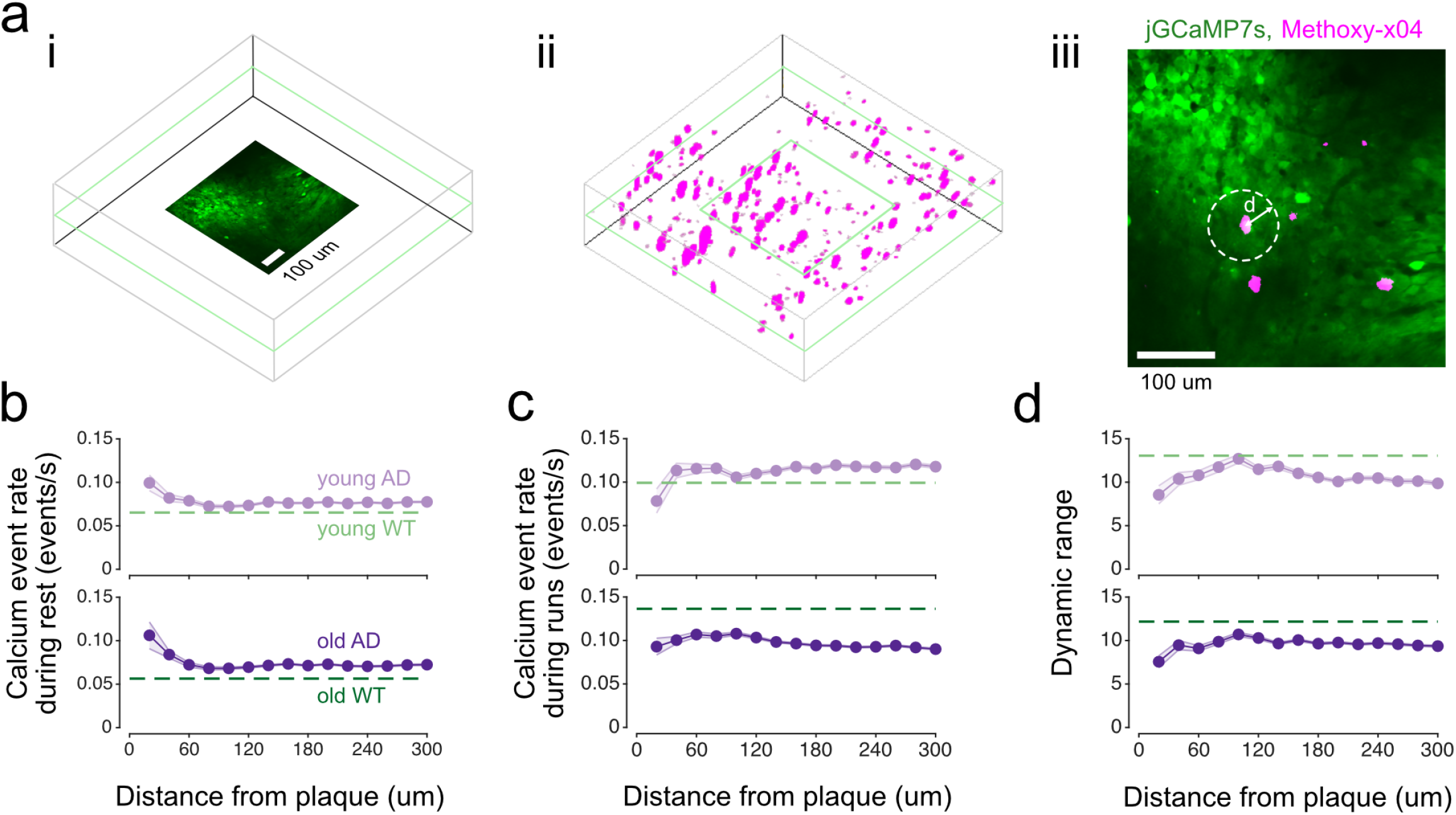
Aberrant excitability and reduced dynamic range of activity near plaques in 5xFAD mice. **a** (i)Three-dimensional volume showing the location of the 490 x 490 *µm*^2^ two-photon jGCaMP7s imaging FOV. (ii) Same 3D field as (i) overlaid with the locations of Methoxy-X04-stained amyloid plaques in a 1100 x 1100 x 300 *µm*^3^ volume relative to jGCaMP7s FOV. (iii) Measurements, e.g. activity rate, are averaged over all cells at a distance *d* from a plaque, shown schematically, then averaged over all plaques. **b** Average neuronal activity versus distance from plaques for young and old 5xFAD mice during rest (*±* SEM). Dashed lines in (b-d) denote the average for the corresponding WT group of the same age. Young 5xFAD: n = 1479 cells pooled from 6 mice; old 5xFAD: n = 1365 cells, 4 mice. For WT means, young WT: 5168 cells, 6 mice; old WT: 3475 cells, 8 mice. **c** Average neuronal activity versus distance from plaques during running. **d** Average dynamic range of neuronal activity during locomotion versus distance from plaques.

In old 5xFAD mice, neural circuit alterations were observed at all plaque distances (Fig. 3b-d). That is, baseline activity was elevated, activity during running was lower, and the speed-related dynamic range was lower relative to the mean of old WT cells regardless of plaque distance. Moreover, the elevation of baseline activity and the reduction in dynamic range was most pronounced near plaques (*<* 20 *µ*m, Fig. 3b,d). In young 5xFAD mice, where neural circuit alterations were not significant at the group level (Fig. 1a, 2b), we also observed elevated baseline activity and reduced dynamic range compared to the mean of young WT cells with the effect being strongest in the vicinity of plaques (*<* 20 *µ*m, Fig. 3b,d). In contrast, calcium event rate during running was elevated in young 5xFAD cells *except* in the vicinity of plaques (*<* 20 *µ*m, Fig. 3b) where it was lower compared to the young WT mean.

Together, these findings suggest that neuronal circuit alterations first emerge near amyloid plaques in young 5xFAD mice and become more widespread with age.

### 2.4 Aberrant neuronal synchrony in 5xFAD mice

Network hypersynchrony in the cortex and hippocampus has been reported in other AD mouse models prior to plaque deposition, through cortical electroencephalogram (EEG) recordings, mapping of chronic hippocampal seizure markers, and resting-state functional magnetic resonance imaging (fMRI) (Verret et al., 2012; Bezzina et al., 2015; Shah et al., 2016). In later stages of the disease, hyposynchrony in resting-state fMRI BOLD signals has also been reported (Shah et al., 2016). We therefore asked if synchrony is altered in the 5xFAD mouse model. To quantify network synchrony, we calculated the Pearson cross-correlation coefficient of neural activity at zero time lag, averaged over all pairs of neurons in each imaging field of view.

During rest, we observed greater synchrony between pairs of cells in young 5xFAD mice compared to that in WT mice (Fig. 4). No significant difference was observed in synchrony in old 5xFAD mice in the resting state. During locomotion, however, the hypersynchrony in young 5xFAD mice disappeared, while significant *hypo*synchrony was observed in old 5xFAD mice. We conclude that there is some evidence for disruption of network dynamics during disease progression in the 5xFAD model.

**Figure 4:**
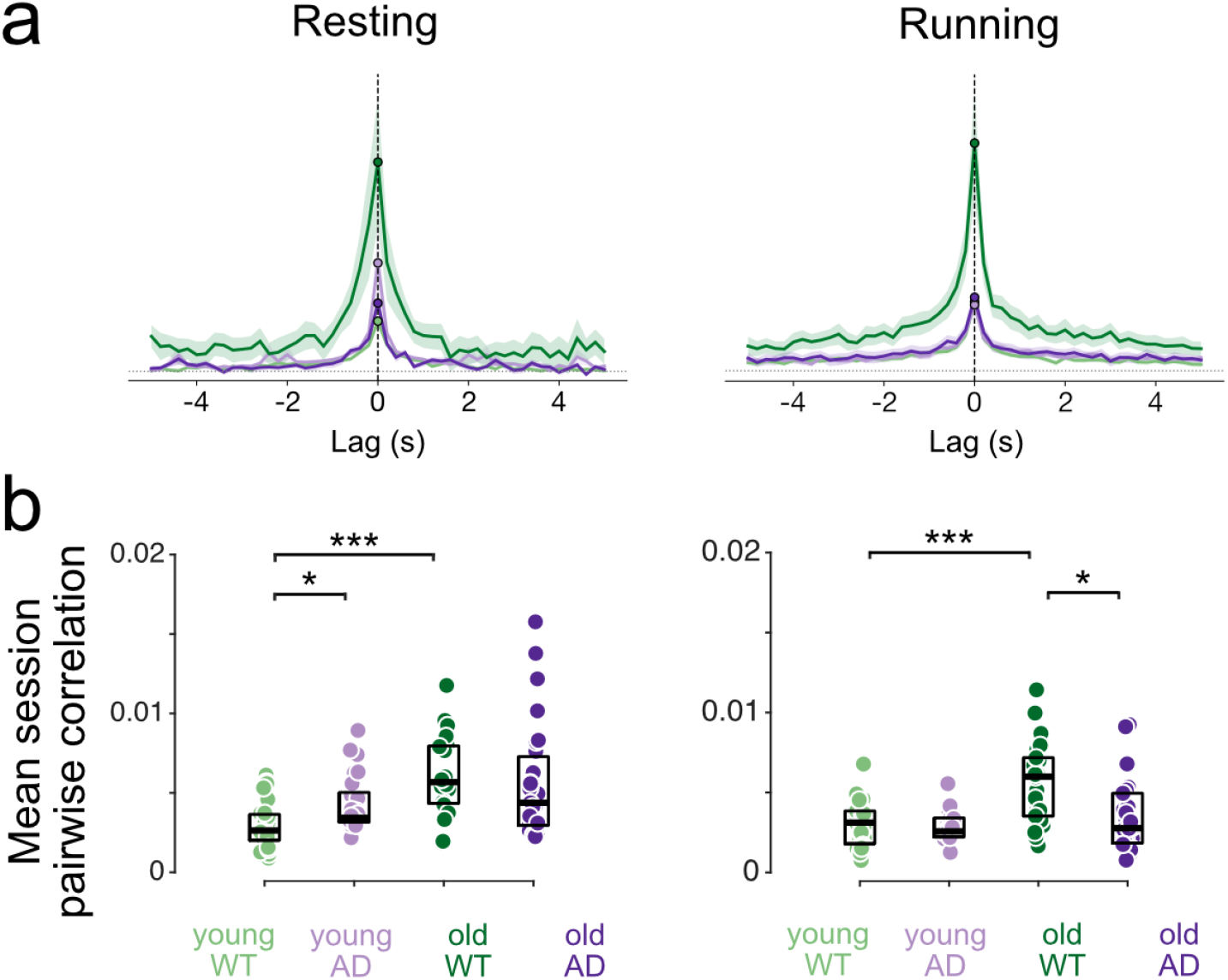
Aberrant synchrony in 5xFAD mice. **a** Mean pairwise cross-correlograms of CA1 event-rate activity during resting (*left*) and running (*right*) periods. Activity was binned in 200 ms windows and cross-correlations were computed over *±* 4s. Curves show group mean *±* SEM across sessions. The vertical dotted line marks zero lag; the horizontal dotted line marks zero correlation. **b** Session-level zero-lag synchrony, measured as the mean pairwise Pearson correlation for different groups during resting (*left*) and running (*right*) periods. Each data point indicates average of all pairwise correlations in an imaging session. Resting, young WT: 0.0029 *±* 0.0002; young 5xFAD: 0.0043 *±* 0.0.0003; old WT: 0.0062 *±* 0.0005; old 5xFAD: 0.0056 *±* 0.0007. Running, young WT: 0.0030 *±* 0.0002; young 5xFAD: 0.0028 *±* 0.0.0002; old WT: 0.0057 *±* 0.0005; old 5xFAD: 0.0035 *±* 0.0004. Statistics were performed using hierarchical bootstrap analysis. For all panels, young WT: n = 5,310 cells pooled from 6 animals, 33 sessions; young 5xFAD: n = 4,955 cells, 6 animals, 27 sessions; old WT: n = 3,517 cells, 8 animals, 27 sessions; old 5xFAD: n = 7,400 cells, 6 animals, 31 sessions. ^∗^*p <* 0.025, ^∗∗∗^*p <* 0.0025. See also Figs. S4 and S5.

### 2.5 Degraded spatial coding in 5xFAD mice

Spatial navigation deficits are an important preclinical marker for AD (Coughlan et al., 2018). Although navigation can be assessed behaviorally, hippocampal CA1 place-cell activity provides a more direct readout of spatial coding that is less dependent on task performance. We therefore quantified spatial information rates in CA1 neurons (Skaggs et al., 1992; Langston et al., 2010). Previous studies have reported that reduced spatial information in hippocampal CA1 cells correlates with plaque load and impaired behavioral performance across mouse models of amyloid pathology (Cacucci et al., 2008; Jun et al., 2020; Chockanathan and Padmanabhan, 2024). However, analyses in the 5xFAD model have produced conflicting results (Zhang et al., 2023; Prince et al., 2021), potentially because of differences in recording modality, such as electrophysiology versus calcium imaging. In our data, spatial information was reduced in old 5xFAD mice compared with old WT mice and young 5xFAD mice (Fig. 5a).

**Figure 5:**
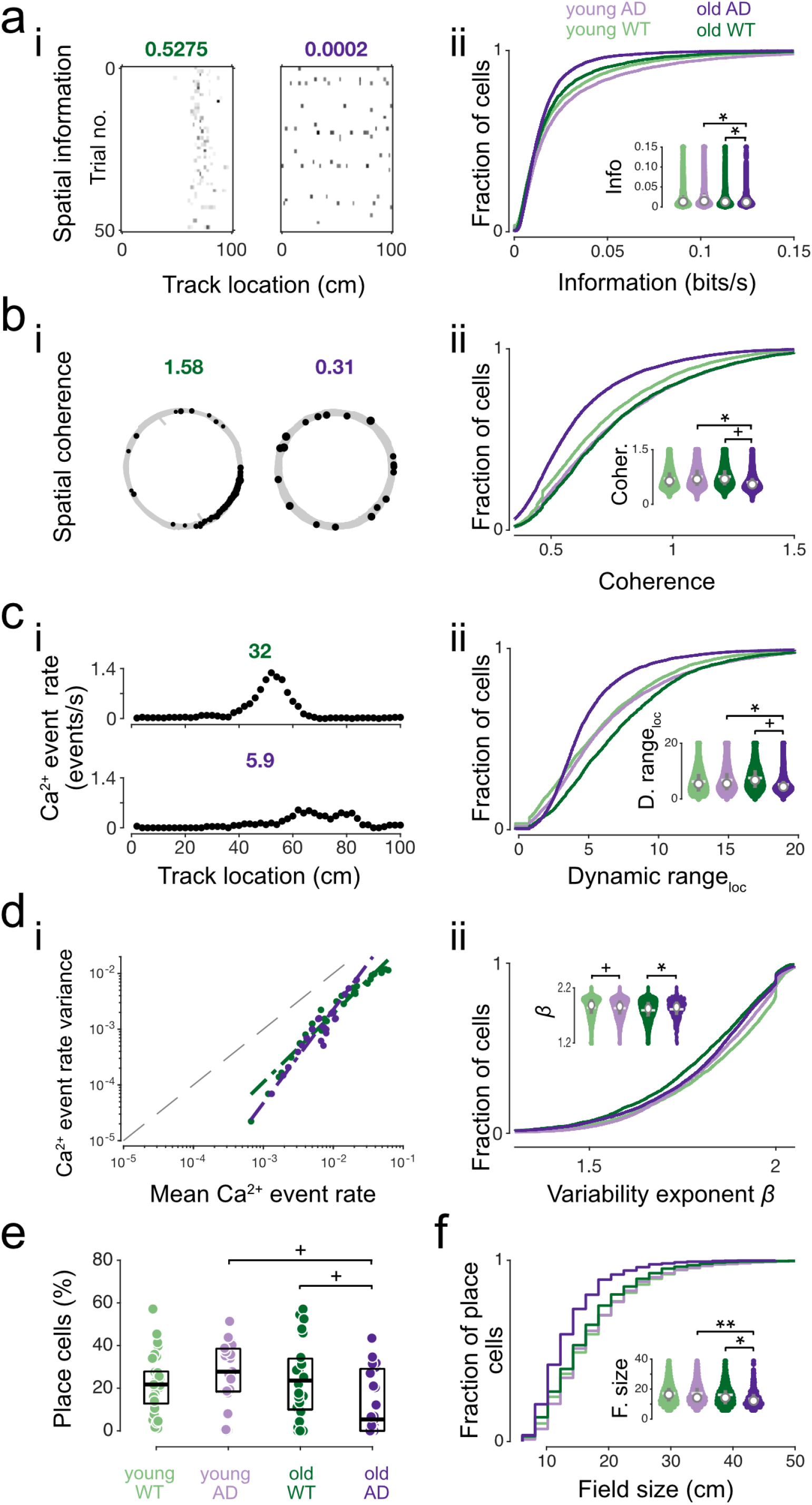
Degraded spatial coding in 5xFAD mice. **a** (i) Neuronal activity maps for two representative cells (one old WT, one old 5xFAD, color codes as in (ii)) with different spatial information values (shown above map). (ii) Cumulative histograms and violin plots (inset) of information comparing WT with 5xFAD for young and old mice. Young WT: 0.023 *±*0.0.002 bits/s; young 5xFAD: 0.029 *±* 0.006 bits/s; old WT: 0.021 *±* 0.002 bits/s; old 5xFAD: 0.017 *±* 0.002 bits/s. **b** (i) Calcium transient location (black dots) overlaid on mouse spatial trajectory (gray line) for two representative cells with different spatial coherence values. (ii) Cumulative histograms and violin plots (inset) comparing average coherence of WT with 5xFAD young and old mice. Young WT: 0.71 *±* 0.02; young 5xFAD: 0.76 *±* 0.04; old WT: 0.76 *±* 0.03; old 5xFAD: 0.60 *±*0.05. **c** (i) Place tuning curves for two representative cells, allowing calculation of spatial response dynamic range (shown above plots), defined as the maximum divided by the non-zero minimum activity across the spatial tuning curve. (ii) Cumulative histograms and violin plots (inset) comparing spatial response dynamic range. Young WT: 6.3 *±* 0.4; young 5xFAD: 6.9 *±* 0.6; old WT: 7.6 *±* 0.4; old 5xFAD: 5.2 *±* 0.6. **d** (i) Variance of calcium event rate in each 2-cm spatial bin plotted against the mean calcium event rate in that bin, for two representative cells. This plot is double-logarithmic, and the slope of the linear fit (total least-squares regression fit, dash-dot line) indicates the exponent (*β*) in the relationship between the variance and mean. In this example, *β* for old WT: 1.23, old 5xFAD: 1.68. (ii) Cumulative histograms and violin plots (inset) of *β* (young WT: 1.82 *±* 0.02; young 5xFAD: 1.83 *±* 0.01; old WT: 1.79 *±* 0.01; old 5xFAD: 1.81 *±* 0.02). **e** Proportion of cells that meet place cell criteria in each group. Young WT: 23.5 *±* 4.2%; young 5xFAD: 26.0 *±* 5.1%; old WT: 29.8 *±*5.5%; old 5xFAD: 11.1 *±* 4.3%. **f** Place field size cumulative histograms (staircase due to 2 cm spatial binning) and violin plots (inset). Young WT: 18.3 *±* 0.8 cm; young 5xFAD: 18.3 *±* 1.1 cm; old WT: 17.8 *±* 0.3 cm; old 5xFAD: 14.4 *±* 0.9 cm. See also Fig. S6. All statistics were performed with hierarchical bootstrap analysis. For A-D, young WT: n = 5082 cells pooled from 6 animals, 31 sessions; young 5xFAD: 4955 cells, 6 animals, 27 sessions; old WT: 3268 cells, 8 animals, 24 sessions; old 5xFAD: 7400 cells, 6 animals, 31 sessions. For E-F, young WT: n = 1470 place cells, 6 animals, 31 sessions; young 5xFAD: n = 1428 place cells, 6 animals, 27 sessions; old WT: n = 1101 place cells, 8 animals, 24 sessions; old 5xFAD: n = 1162 place cells, 6 animals, 31 sessions. ^+^*p <* 0.05, ∗*p <* 0.001, ∗ ∗ *p <* 0.005.

To obtain a broader assessment of spatial coding quality, we next examined complementary measures of spatial tuning. Spatial coherence, which quantifies how spatially contiguous a neuron’s activity is, was similar in young WT, young 5xFAD, and old WT mice, but was significantly lower in old 5xFAD mice (Fig. 5b). We also measured the dynamic range of place tuning, defined as the ratio between the maximum and non-zero minimum values of the place tuning curve, which quantifies the capability of a neuron to discriminate location. This measure was similar in both groups of young mice, but was significantly reduced in old 5xFAD mice (Fig. 5c).

We then asked whether trial-to-trial response variability was altered in 5xFAD mice. Reliable spatial coding requires neurons to respond consistently when the animal revisits the same location. However, neural responses typically vary across repeated presentations of the same stimulus or behavioural condition, and this variability often increases with mean activity (Tolhurst et al., 1983). We previously observed a similar relationship in CA1 neurons (Go et al., 2021). Because response variability can reflect both local amplification and network dynamics (Carandini, 2004), we reasoned that it might be sensitive to AD-related circuit disruption.

For each cell, we plotted the variance of calcium event rate over laps against the mean event rate in each spatial bin (Fig. 5d i; see Go et al. (2021) for a more detailed description of this analysis approach). We quantified this relationship using the slope or variability exponent of the fitted line in log-log coordinates. A higher slope indicates that neuronal response variability scales more rapidly with event rate, which we have proposed (Go et al., 2021) to be a measure of intrinsic cellular variability and may be sensitive to disruptions in the local synaptic and cellular environment. In our analyses, old 5xFAD mice showed increased variability slope compared with old WT mice (Fig. 5d-ii). Young 5xFAD mice also showed higher variability exponents compared to young WT mice.

Finally, we assessed place-cell prevalence and field size. Old 5xFAD mice had a smaller fraction of place cells than old WT mice (Fig. 5e). Their place fields were also smaller (Fig. 5f). Although this finding of reduced field size differs from reports in some other AD mouse models (Takamura et al., 2021; Jun et al., 2020), it is consistent with previous observations in the 5xFAD model (Zhang et al., 2023).

Together, these complementary measures indicate degraded spatial coding in old 5xFAD mice, including reduced spatial information, lower spatial coherence, reduced place-tuning dynamic range, increased trial-to-trial variability, fewer place cells, and smaller place fields. We next examined whether these spatial coding deficits depend on the 3D proximity of neurons to amyloid plaques. We found a plaque-proximity effect in several measures of spatial coding in old AD mice: neurons closer to plaques generally exhibit poorer spatial tuning, with lower spatial information, higher variability and smaller field size (Fig. S6a,d-f). These distance-dependent changes tend to diminish with increasing distance from plaques. Coherence and place-tuning dynamic range are lower in old AD than old WT across essentially the full range of plaque distances. In young AD mice, however, these measures are higher than in young WT mice except close to plaques where they are lower, suggesting that disruption begins near plaque locations. Overall, across multiple spatial coding measures, the common trend is that plaque-adjacent neurons in old AD mice display the greatest disruption.

### 2.6 Delayed place tuning in 5xFAD mice

To test whether AD impacts encoding and recall differently, we imaged mice first in a familiar track (FAM) and then a novel track (NOV) within the same recording session. The familiar track was the training environment while the novel track was introduced only on the imaging day. The two tracks differed in visual and tactile cues on the walls and floor. Only sessions for which mice ran 40-50 laps in each environment were included in the analysis.

We first assessed within-session place field stability by dividing each session into two equal halves and calculating the correlation between the place fields derived from each half. In young mice, place fields were similarly stable in WT and AD animals in the familiar environment, but were less stable in both groups in the novel environment (Fig. 6a). In old mice, however, AD place fields were less stable than WT place fields in the familiar environment, and stability declined further in the novel environment, where old AD mice showed the lowest stability overall (Fig. 6a).

**Figure 6:**
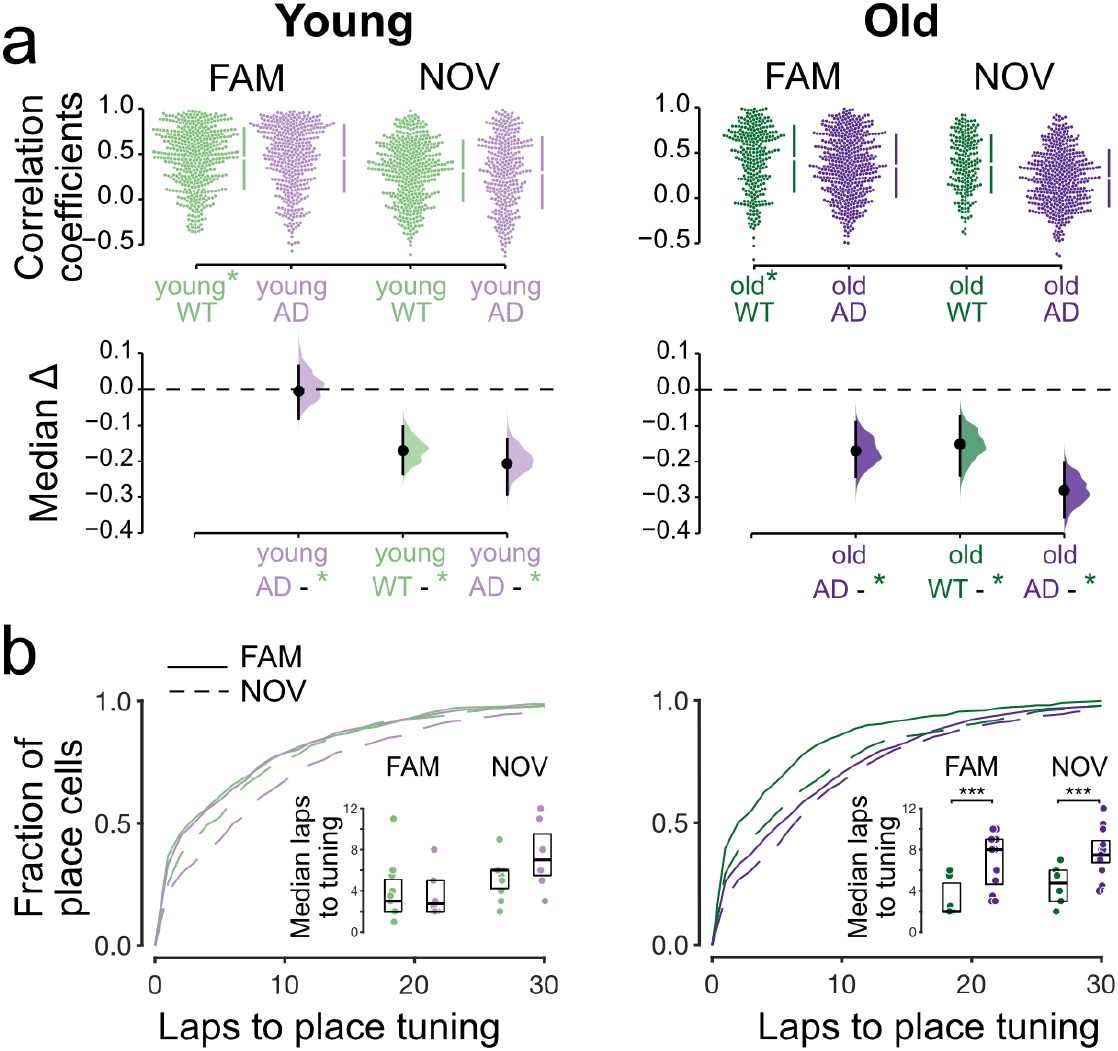
Delayed tuning of place fields in 5xFAD mice. **a** Intra-session stability of place fields, quantified by the correlation between the place fields of two halves of the trial. Data shown for familiar (FAM) and novel (NOV) environments. Cumming plots are used to visualize the bootstrap effect size. Top panels show distribution of correlation coefficients, bars indicate mean and standard deviation; bottom panels show median difference with 95% confidence interval, with left-most group as reference. **b** Cumulative histograms of number of laps before place tuning is reached for the different groups. (*Insets*) Median number of laps to place tuning. Each data point corresponds to a session for one mouse. (*Left*) FAM, young WT: 3.8 *±* 0.3 laps; young 5xFAD: 3.7 *±* 0.3 laps. NOV, young WT: 5.3 *±* 0.2 laps; young 5xFAD: 7.4 *±* 0.4 laps. (*Right*) FAM, old WT: 3.1 *±* 0.2 laps; old 5xFAD: 6.9 *±* 0.3 laps. NOV, old WT: 4.6 *±* 0.3 laps; old 5xFAD: 7.7 *±* 0.2 laps. \*\*\**p <* 0.002, hierarchical bootstrap test. See also Fig. S7. For all panels, young WT: n = 449 place cells pooled from 6 animals, 11 sessions; young 5xFAD: n = 390 place cells, 5 animals, 8 sessions; old WT: n = 280 place cells, 6 animals, 6 sessions; old 5xFAD: n = 291 place cells, 5 animals, 8 sessions.

We next examined when place tuning emerged within each session by identifying the first lap in which each place cell showed place tuning. In all groups, some cells were tuned from the start of the session, whereas others developed tuning gradually with experience (Fig. 6b).

Our first hypothesis was that Alzheimer’s disease impairs recall in the familiar environment, leading to reduced within-session place field stability. Consistent with this idea, place cells in old AD mice required more laps to become tuned in the familiar track than those in old WT mice (Fig. 6b, right inset). By contrast, young AD mice did not differ significantly from young WT mice (Fig. 6b, left inset).

Our second hypothesis was that Alzheimer’s disease slows the encoding of new place fields, thereby reducing intra-session stability in the novel environment. This was also supported by the data: in the novel track, place cells in old AD mice required more laps to become tuned than those in old WT mice, indicating slower encoding (Fig. 6b, right inset). Young 5xFAD place cells showed only a modest, non-significant increase in tuning latency relative to young WT cells, suggesting a weaker effect at this age (Fig. 6b, left inset).

To better understand the basis of these delays, we divided place cells into those tuned at session onset and those that became tuned during the session. In the familiar environment, old AD mice had fewer cells tuned from the start than age-matched WT mice (Fig. S7a), consistent with impaired recall. In the novel environment, old AD mice also had fewer initially tuned cells (Fig. S7a), consistent with slower encoding. In both environments, old AD mice showed slower tuning acquisition among cells that were not initially tuned (Fig. S7b). Young AD mice did not differ from young WT mice in the initial proportion of tuned cells, but in the novel track they were slower to acquire tuning after the initial lap (Fig. S7).

Overall, these results indicate that place tuning emerges more slowly in 5xFAD mice, especially in older animals. The effect is evident in both familiar and novel environments, suggesting impairments in both recall of established spatial representations and encoding of new ones.

## 3. Discussion

Our main finding is that amyloid pathology in 5xFAD mice alters CA1 activity dynamics by increasing baseline activity while reducing the increase in firing that is normally expected during locomotion. Together, these changes reduce the dynamic range of neuronal activity. To our knowledge, this is the first direct experimental evidence for reduced dynamic range in an AD mouse model. We propose that this reduction is functionally important because it fundamentally limits the capacity of neurons to modulate their responses in response to behavioural states, thereby degrading information coding in hippocampal circuits. This reduction in dynamic range was accompanied by deficits in both speed coding and spatial coding, as well as slower place-field emergence in familiar and novel environments. These findings suggest that amyloid pathology impairs not only basal excitability, but also the flexible modulation of CA1 activity required for accurate behavioral and spatial representations.

The elevated baseline activity we observed in old 5xFAD mice is consistent with earlier reports in anaesthetised mice (Busche et al., 2012; Šišková et al., 2014). Busche et al. (2012) described a complex excitability scenario in the APP/PS1 mouse, with a marked increase in the fraction of both “silent” (or hypoactive) neurons and hyperactive neurons in the plaque-bearing CA1 region of older transgenic mice. We see a similar trend in our data. In older AD mice, there is a marked increase in the number of cells with near-zero activity rates (Fig. 1d,e).

In behaving mice, there have been conflicting reports of increased, decreased or no difference in activity in AD mice (Takamura et al., 2021; Mably et al., 2017; Zhang et al., 2023; Jun et al., 2020; Lin et al., 2022) either in resting or running states. This discrepancy may be due to different mouse models, different measurement modalities (electrophysiology versus calcium imaging; in calcium imaging, further difference may stem from whether or not amplitudes of individual events are accounted for), but may be largely due to different speed thresholds (ranging from *>* 0 to *>* 3 cm/s) for defining the mobile state. We speculate that our conservative choice of speed thresholds for the resting (*<* 1 cm/s) and running (*>* 2 cm/s) states may reconcile the discordant results in literature.

We observed higher average running speeds in our head-fixed old 5xFAD mice as they traversed a circular track. This hyperactivity in behavior was not observed in APP knock-in mice (Takamura et al., 2021) or in tethered 5xFAD mice exploring an open field (Zhang et al., 2023). It has however been observed in APP-PS1 mice (Lagadeca et al., 2012) and is consistent with restlessness which has been noted in AD patients (Teri et al., 1988). Because locomotor behavior differed across groups, analyses were restricted to well-sampled speed ranges (Fig. S2b), reducing the influence of unequal occupancy across speed bins.

A key advantage of our two-photon approach is that it enabled us to relate neuronal functional changes to plaque proximity. In young 5xFAD mice, abnormalities were largely confined to neurons near plaques. In old 5xFAD mice, however, the same alterations were more spatially widespread, although still strongest close to plaques. This pattern suggests a progression in which plaques act as local foci of dysfunction early in disease, followed by broader network disruption with age. The local effects we observed are consistent with previous work showing that amyloid plaques are associated with aberrant activity (Busche et al., 2008) and dendritic structural disruption (Le et al., 2001) in nearby neurons, as well as evidence that dendritic degeneration is linked to increased excitability (Šišková et al., 2014).

Dendritic degeneration may possibly explain the reduced dynamic range in old 5xFAD mice. Dynamic range has been linked to dendritic architecture, with larger and more complex dendritic trees supporting broader response ranges (Gollo et al., 2009). Given the well-established dendritic spine loss and arbor degeneration in AD (Moolman et al., 2004; Tsai et al., 2004; Grutzendler et al., 2007; Šišková et al., 2014), it is plausible that dendritic pathology contributes to the reduced dynamic range observed here. As a larger dynamic range is believed to increase the probability of a neuron’s survival (Gollo et al., 2012; Wang et al., 2017), a reduced dynamic range may be particularly relevant in the context of neurodegeneration characteristic of AD. Reduced dynamic range may also relate to disrupted gamma oscillations in AD. Gamma rhythms support hippocampal coding and ensemble coordination. Altered gamma activity in 5xFAD mice (Iaccarino et al., 2016) may therefore contribute to a reduced ability of CA1 circuits to modulate firing across behavioral states.

The functional consequences of activity changes in old 5xFAD mice were evident in hippocampal coding. Old 5xFAD mice showed poorer spatial coding across several measures, including reduced spatial information, lower spatial coherence, reduced place-tuning dynamic range, increased trial-to-trial variability and fewer place cells. This is consistent with earlier reports of impaired spatial representation in AD (Cheng and Ji, 2013; Mably et al., 2017; Jun et al., 2020). The increase in response variability may be especially important. Reliable spatial coding requires neurons to respond consistently across repeated traversals of the same location. Increased variability reduces the fidelity of neural representations and provides one possible mechanism for the lower spatial information and weaker place coding we observed.

The spatial coding deficits were again most pronounced near plaques, indicating that local amyloid pathology is linked not only to altered excitability, but also to degraded spatial representations. In young 5xFAD mice, some spatial coding measures were altered primarily near plaques, consistent with an earlier, more localized stage of impairment.

We also found evidence that Alzheimer-like pathology impairs both recall of established spatial representations and encoding of new ones. In the familiar environment, place fields in old 5xFAD mice were less stable and emerged more slowly (Fig. 6a), suggesting impaired recall of an existing spatial map. In the novel environment, place tuning also developed more slowly, consistent with impaired encoding of new spatial representations. These delays were strongest in old 5xFAD mice, although young 5xFAD mice also showed weaker effects in some analyses. Together, these results indicate that amyloid pathology affects multiple stages of hippocampal spatial processing, from the stability of existing maps to the formation of new ones.

Our findings are consistent with studies reporting impaired memory recall in AD animal models (Zhao et al., 2014; Poll et al., 2020) and in human AD patients (Haj and Robin, 2021), and with reports of limited refinement in spatial tuning in animals learning to navigate a novel environment (Zhao et al., 2014; Broussard et al., 2022). Reduced sharp wave ripples (SWRs) observed in both young (Iaccarino et al., 2016) and old (Prince et al., 2021) 5xFAD mice may contribute to the slower emergence of place fields observed here. Because SWRs support replay and place field consolidation, weaker ripple activity could impair the stabilization of both familiar and newly forming spatial representations.

At the network level, coordination was also altered. During rest, young 5xFAD mice showed increased synchrony, whereas during locomotion old 5xFAD mice showed reduced synchrony. These results suggest that amyloid pathology does not shift synchrony uniformly in one direction, but instead alters coordination in a state- and stage-dependent manner. One possibility is that elevated baseline activity promotes excessive co-activation at rest early in disease, while later degeneration and degraded coding reduce coordinated activity during behavior. Interneuron dysfunction may provide a circuit-level explanation for these effects. Because inhibitory interneurons regulate excitability, gain, and rhythmic synchronization, their disruption could compress dynamic range and alter both gamma-related coordination during behavior and ripple-related coordination during rest. Because coordinated hippocampal activity is important for memory consolidation, these synchrony changes may contribute to the learning deficits we observed.

The old group spans 6.4–10.1 months (median 7.5), a range over which 5xFAD pathology progresses substantially. Because this heterogeneity could mask or inflate effects, we re-analyzed our results excluding animals with age 9 months old and above and found that our main results remain unchanged. This shows that our findings are not driven by the oldest animals.

In summary, 5xFAD mice showed a progression of hippocampal dysfunction that began locally near plaques in young (2.7 month old median age) animals and became more widespread in older (7.5 mo) animals. The major features of this dysfunction were elevated baseline activity together with reduced locomotion-driven firing leading to a reduction in dynamic range; altered synchrony, degraded spatial coding, increased response variability, and delayed acquisition of spatial tuning in both familiar and novel environments, indicative of deficits in both recall and learning. Our work offers new insights into the progression of neural circuit abnormalities in mouse models of amyloid disorders, and how these effects lead to memory and cognitive impairment, with potential implications for the development of new therapeutic approaches for AD.

## 4. Acknowledgements

This work was supported by UKRI/Wellcome grant EP/W024020/1 to SRS; Alzheimer’s Research UK (ARUK-NC2019-IMP) grant to SRS and MAG; EPSRC grant EP/J021199/1 to SRS; EPSRC CDT in Neurotechnology for Life and Health (EP/L016737/1) studentship to SVP; BBSRC grant BB/R022437/1 to SRS; Wellcome Trust grant 221522/Z/20/Z to SRS; Chan-Zuckerberg Initiative award NC/W000903/1 to SRS, and a philanthropic donation from Mrs Anne Uren and the Michael Uren Foundation to SRS. We thank M. Sastre for providing the 5xFAD mice, and D. Dupret, N. Zabouri, R. Mitchell-Heggs, M. Sastre and S. Barnes for useful discussions.

## 5. Rights retention

This research was funded in whole, or in part, by the Wellcome Trust [Grant number 221522/Z/20/Z]. For the purpose of open access, the author has applied a CC BY public copyright licence to any Author Accepted Manuscript version arising from this submission.

## Contributions

MAG and SRS conceived the study and designed the experiments; MAG and YL performed the imaging and behavioural experiments; MAG, KEC, SVP, BRFT, MG, JJY and SRS analyzed the data; MAG, KEC and SRS wrote the article.

## Methods

### Ethical approval declarations

All experimental procedures were carried out under the Animals (Scientific Procedures) Act 1986 and according to Home Office and institutional guidelines.

### Animals

Male hemizygous 5xFAD mice (B6SJL-Tg(APPSwFlLon,PSEN1*M146L*L286V)6799Vas/Mmjax, MMRRC stock # 034840-JAX) were crossed with female C57BL6/J mice (JAX stock # 000664) to maintain 5xFAD and non-transgenic wild-type colonies under standard animal breeding at Imperial College London. Non-transgenic littermates were used as controls. The animal room was kept on a 12:12 light-dark cycle daily. Animals were divided into two age groups: young group aged 1.5-2.4 months at the time of viral injection, 2.1-3.0 months at the time of imaging; and old group aged 5.6-9.3 months during viral injection, 6.4-10.1 months during imaging. Data were analyzed from 6 young wild type (2 females, 4 males), 6 young 5xFAD (3 females, 3 males), 8 old wild type (5 females, 3 males) and 6 old 5xFAD mice (3 females, 3 males). Both male and female mice were used for this study. Animal sample sizes were determined by availability; all animals for which successful imaging was carried out were analyzed.

### Experimental Methods

#### Virus injection and hippocampal window surgery

Surgical procedures followed Go et al. (2021) closely. Mice were anaesthetised with 1.5-3% isofluorane (Iso-Vet, Chanelle Pharma) and body temperature was maintained at 37^°^C. Analgesia was administered pre-operatively with Carprofen (5 mg/kg, Rimadyl^®^, Zoetis) and buprenorphine (0.07 mg/kg, Vetergesic^®^, Ceva Animal Health Ltd). Anaesthetic depth was assessed via pedal flexion every 10 min for the duration of surgery. A small (∼0.5 mm) craniotomy was made and a cocktail of two viruses (pGP-AAV-syn-jGCaMP7s-WPRE, Addgene catalog 104487-AAV9, titer 1.9 1*×* 013 vg/ml, ∼50 nL and pAAV-CAG-tdTomato, Addgene catalog 59462-AAV1, titer 1.9 *×* 1013 vg/ml, ∼50 nL) was injected into the hippocampal CA1 region (coordinates from bregma, in mm: -1.3 and -1.5 DV, -1.8 ML, 2.0 AP). The first virus delivers a green genetically encoded calcium indicator (jGCaMP7s), and the second, an activity-independent red fluorescent protein (tdTomato) which is helpful for tracking cell body movements over time. pAAV1-hSyn1-mRuby2-GSG-P2A-GCaMP6s-WPRE-pA, which expresses the red fluorescent (mRuby2) and the green calcium indicator (GCaMP6s), was injected in a few early cohorts of animals. Two weeks post-injection, a hippocampal window was implanted as described by Dombeck et al. (2010). A circular craniotomy was made and the cortex (including parietal cortex and parts of visual and hindlimb sensory cortex) above the injection site was aspirated using a 27 gauge needle connected to a water pump until the fibers of the corpus callosum became visible. A stainless steel cannula (diameter: 3 mm, height: 1.5 mm) with a glass bottom was pressed down into the tissue and fixed in place using histoacryl glue (B.Braun Surgical). The surrounding skull was roughened using a scalpel blade and a stainless steel headplate (aperture: 8.5 mm) was glued to the skull, centred on the craniotomy. Exposed skull outside the headplate aperture was covered with dental cement mixed with black powder paint. Mice were given carprofen (5mg/kg/24hrs) in oral water for 3 days after surgeries. Mice were allowed 5-7 days to recover before behavioural training began.

#### Behavioral training

Behavioral training procedures are described in detail in Go et al. (2021). Approximately one week after the hippocampal window was implanted, the animals were habituated to the apparatus and experimenter, and placed under water restriction. Animals were trained to move in the dark along a circular track (outer diameter: 32.5 cm, width: 5 cm) floating on an air table (Mobile HomeCage Large, Neurotar). Infrared (IR) light illuminated the training area and an IR camera was used to monitor the animals. The floating tracks were made of carbon fibre (weight: 100 *±* 2.8 g) and had 4-cm high walls lined with visual (phosphorescent tape, Gebildet E055 and E068) and tactile cues (sandpaper, cardboard, foam, bubble wrap). The phosphorescent tapes emitted light at 500 (blue) and 520 (green) nm throughout the imaging session (Go et al., 2021). Mouse position on the track was measured using a magnetic tracking system (Neurotar) which enabled closed-loop position-based reward delivery via a lickspout. Liquid rewards were accompanied by a beep.

Animals were trained twice daily in 45-min sessions. Mice were trained with one circular track in the morning and a second circular track with different visual and tactile cues in the afternoon. Water rewards of 4 *µ*L per loop traversed were delivered at random locations. Daily water intake was limited to 1-3 mL individually adjusted for each mouse to maintain 85% of pre-restriction weight. At the start of each training session, two rewards were given to motivate the mice to lick for water. There was no limit to the number of rewards animals could receive during sessions. If the animal did not reach the target volume for the day during training, the remaining volume was given at the end of the last training session for the day. Animals were trained for 11-14 sessions before imaging sessions began.

#### In vivo two-photon imaging

Imaging was performed with a two-photon resonant scanning microscope (Scientifica VivoScope) equipped with a tiltable objective mount and a 16 *×* water-immersion objective (LWD 0.8 NA, Nikon). We used 50% concentration ultrasound gel as the immersion fluid. For imaging jGCaMP7s and tdTomato, laser (Mai Tai HP, Newport) wavelength was set to 940 nm. Laser power underneath the objective was 60 - 166 mW. Mice were imaged while navigating a familiar (floating) circular track in sessions which lasted up to an hour. Images (512 *×* 512 pixels, 490 *×* 490 *µ*m field of view) were acquired at 30 Hz. In learning experiments, mice were imaged for up to an hour in a familiar track then moved to a novel track where they were imaged for up to another hour. Sessions for which mice ran 40-50 laps were included for analysis. SciScan software (Scientifica) was used for microscope control, and image acquisition was TTL-synchronized to position tracking and reward timing signals. Mouse position and speed data were acquired at 100 Hz with the Neurotar software. Light from the phosphorescent tapes on the floating track walls used as visual cues was not detectable on either red or green imaging channels.

After the last imaging session, amyloid plaques were mapped *in vivo* at least 24 hours following i.p. injection of Methoxy-x04 (dose: 10 mg/kg), using 740 nm laser excitation.

#### Calcium imaging data processing

Processing of two-photon calcium imaging data was performed as described in Go et al. (2021). A customised image processing pipeline based on the MATLAB version of CaImAn (Giovannucci et al., 2019) was used. Motion artefacts were removed using rigid and then non-rigid image registration. Regions of interest (ROIs) were automatically identified and overlapping ROIs excluded. To track cells across multiple images, we temporally concatenated the images and segmented the concatenated video. Neural activity was deconvolved from the fluorescence traces using OASIS (Friedrich et al., 2017).

### Data Analysis

All analyses were performed with custom MATLAB scripts.

#### Temporal binning

We used temporally binned activity to reduce calcium-event timing uncertainty. The neural activity temporal resolution is defined by the image sampling rate (30 Hz). We reduced the temporal resolution of neural activity by summing event counts in 6 consecutive time bins (effective rate 5 Hz). For speed and position data, we first downsampled the data from 100 Hz to 30 Hz. We then took the average value over 6 time bins to match the temporal resolution of the neural activity. No temporal binning was done in the calculation of activity rate variability.

#### Neuronal activity rate

For the analyses in this paper, except for spatial information calculation, activity rate was calculated by integrating the number of calcium events within a 200 ms time window and dividing by window duration and is reported in events per second. For spatial information, activity rate was calculated by integrating the amplitude of events within a 200 ms time window and dividing by window duration.

#### Resting and running periods

Periods in which animal speed was less than 1.0 cm/s were considered resting periods. Periods in which animal speed exceeded 2.0 cm/s were considered running periods. Analyses on speed modulated cells, and place tuned cells were limited to running periods to distinguish speed and place-correlated effects from those related to changes in animal behavior (grooming, pausing for rewards, foraging).

#### Speed-related dynamic range and speed modulated cells

We used 1 cm/s speed bins and calculated speed tuning curves for each cell by dividing the total activity for a speed bin by the total time spent in it, then smoothing using a boxcar average over 3 bins. The dynamic range of activity due to locomotion was defined as the ratio of the maximum of the speed tuning curve and the non-zero minimum.

Following Kropff et al. (2015), neurons were classified as speed modulated cells if their speed response modulation depth exceeded chance level. The speed response modulation depth of a cell was defined as the maximum of its speed tuning curve minus the non-zero minimum. Chance-level was determined by a shuffling procedure. For each shuffle, the neural activity time series was time-shifted by a random number between 10 seconds and the time series duration minus 10 seconds. A speed tuning curve and its corresponding modulation depth was then calculated for the shuffled neural activity. The cell was considered speed modulated if its modulation depth exceeded the 95th percentile of the distribution of values collected from 1000 shuffles.

Following Muzzu et al. (2018), we classified speed modulated cells according to their responses as positively modulated, negatively modulated or having a preferred speed. For each speed modulated cell, we took the running segment of the speed tuning curve (*>* 2 cm/s), included speed bins with total duration of at least 4 seconds, then used the best-fit curve (linear, quadratic, double exponential, Gaussian, or double Gaussian) to determine the response type of the cell. Specifically, speed modulated cells were classified as 1) positively modulated if their maximum firing rate exceeded the firing rate during rest (*<*0.5 cm/s) and was recorded at a speed greater than 70% of the maximum animal speed; (2) negatively modulated if their minimum firing rate was less than the firing rate during rest and was recorded at a speed greater than 70% of the maximum animal speed; (3) having a preferred speed if the maximum firing rate exceeded the firing rate during rest and was recorded at a speed less than 70% of the maximum animal speed.

#### Amyloid plaque detection

Two-photon images of amyloid plaques were averaged for each *z* plane and denoised with a gaussian filter. Images from adjacent FOVs were stitched to create a montage 1.1 um x 1.1 um centred on the imaging FOV. We developed a convolution neural network (CNN) for automatic detection of the centroids of amyloid plaques in images. The training data set consisted of 70 images with manually selected plaque ROIs, augmented to 3360 images by one or more of the following processes: vertical and horizontal flipping, rotations of 90^°^, 180^°^ and 270^°^, and geometrical operations of rotation, zoom, skew, random distortion and shear. The samples were randomly split into 75% for training and 25% as testing data for validation.

#### Analysis of distance to plaques

To examine the spatial relationship between amyloid plaques and cells, the distance from each cell to every plaque was measured. For each plaque, annular rings of width 20 *µ*m were drawn centred on the plaque, with the outer radius (i.e. distance from a plaque) ranging from 20 to 200 *µ*m. Cells were included in the annular ring where they were located and a cell could be included in multiple annular rings of different outer radii centred on different plaques. The quantity of interest (e.g. activity rate, dynamic range, spatial information, etc.) was averaged over all cells within all annular rings of the same outer radius.

#### Synchrony

We calculated the Pearson correlation coefficient for all cell pairs that were imaged in a field of view, normalized such that the auto-correlations at zero lag were identically 1.0. Synchrony was defined as the mean pairwise correlation at zero time lag. Averages were calculated by first taking the Fisher’s transform of the Pearson’s *r* values: *z* = 0.5 ln ((1 + *r*)*/*(1 −*r*)), calculating the mean of the *z* values, and then transforming the average *z* value back to an *r* value.

#### Spatial coding analysis

Mouse position was linearized by converting angular distance to Euclidean distance using the known circumference of the circular track. Neural activity and position time series were sampled at 200 ms bins. We divided the track into 2-cm spatial bins and constructed neural activity rate maps for each cell by dividing the total calcium event count during the occupancy of a spatial bin by the total time it was occupied, smoothing using a boxcar average over 4 bins and normalizing each map by its maximum value.

#### Spatial information

Spatial information rate (in bits/s) for each cell was computed as described previously (Skaggs et al., 1992, 1996):

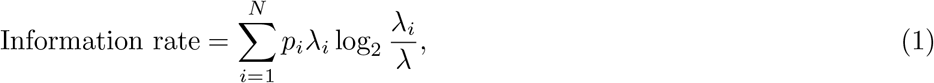

where *i* indexes over *N* spatial bins, *p*_*i*_ is the probability of the animal being in spatial bin *i, λ*_*i*_ is the mean activity rate for bin *i*, and *λ* is the overall mean activity rate of the cell. Activity rate was calculated by integrating the total amplitude of events within a 200 ms time window and dividing by the window duration. Spatial information rate may alternatively be calculated from two-photon calcium imaging data using binarized calcium event rates alone (by analogy to spikes), however each calcium transient event may reflect multiple action potentials, and thus using the amplitude rate is a more accurate (and potentially more sensitive) information measure.

#### Coherence

Spatial coherence is a measure of the spatial contiguity of the activity of a neuron. Similar to Zhang et al. (2014), it was obtained by calculating the correlation coefficient between a neuron’s activity rate map and the same rate map smoothed by a boxcar average over 4 bins. Coherence was defined as the Fisher’s transform of the Pearson’s *r* value: *z* = 0.5 ln((1 + *r*)*/*(1 − *r*)).

#### Spatial response dynamic range

The dynamic range of activity in response to location was defined as the maximum of the activity rate map divided by the non-zero minimum.

#### Neuronal variability analysis

Neuronal variability analysis was performed by adapting methods from classical visual neuroscience literature (Tolhurst et al., 1983) to hippocampal place cells and calcium transient event trains, similar to Go et al. (2021). This analysis shows how variable responses are to laps around the same track. Briefly, for each cell, we calculated for each lap the calcium event rate for each 2-cm bin in the track. We then calculated the mean and the variance of this quantity across laps for all spatial bins for each cell. We used total least squares linear regression to fit a power law model *y* = *αx*^*β*^ to the relationship between activity variance and mean for each individual cell. The vertical intercept parameter *α* describes the overall variability of the cell, with a Poisson process having *α* = 1. Of interest is the power law exponent *β*, which can be read off from the slope of the fitted line, describing how the variability scales with firing rate. We suggest that the variability exponent provides potentially highly useful information about the reliability of neuronal signaling - with disturbances of excitability altering the scaling behavior. An exponent of 1 indicates reliability equivalent to that of a Poisson process; above 1 implies additional sources of variability.

#### Place cell classification

Neurons were classified as place cells if they met the following criteria: (1) calcium transient events were present for at least 60% of the laps through the circular track and (2) the cell contained spatial information greater than chance. Chance-level spatial information for a cell was determined by a shuffling procedure. For each shuffle, the neural activity time series is time-shifted by a random number between 10 seconds and the time series duration minus 10 seconds. The spatial information rate for the shuffled neural activity is then calculated and chance-level values are pooled from 1000 repetitions. The cell was considered a place cell if its information rate exceeded the 95% percentile of the shuffled data. The location of the place field was defined by the bin location of maximum activity rate while the place field size was determined by the number of bins for which the neural activity rate was at least 50% of the maximum.

#### Place field stability

To quantify within-session place field stability, each session was divided into 2 halves. For each cell, the Pearson correlation coefficient between activity rate maps generated from each segment was then calculated.

#### Place field tuning acquisition

For each place cell, the lap of the track at which place cell tuning begins was determined using the binary place field locations identified from the first 50 laps. We looked at firing rates in the place field in a rolling 5 lap window, to find the first tuned 5 lap window. We defined being tuned as a window where the firing rate in the place field locations exceeded the average firing rate from all locations. We assessed this on the whole 5 lap window, as well as for the individual laps. The first window in which the place field firing rate was greater than the mean on more than one lap as well as the whole window was taken to be the tuning acquisition lap. Cells that met the above criteria at no point in the first 50 laps were removed from analysis.

To compare between groups we calculated the median tuning acquisition lap for each recording sessions. Additionally, we measured the proportion of place cells where tuning was present on the first lap in each recording session. Finally, we compared the median tuning acquisition lap of cells where tuning was not present on the first lap. These statistics were determined using session averages pooled over all sessions in each group.

### Statistics and Reproducibility

#### Blinding

As amyloid plaques were visible in the brains of 5xFAD mice, it was not possible to perform blinding at the point of data collection. However, analysis was performed by automated batch scripts, such that blinding effectively occurred in data analysis up to the point of final figure output.

#### Hierarchical bootstrap

In this paper, we consider many neurons which are hierarchically grouped, and as such, not all neurons in a group may be independent samples. Summary statistics (e.g. mean) reported here are obtained using the pooled distribution of all neurons, but a bootstrapping approach (Saravanan et al., 2020) is used to account for the dependence inherent in the hierarchical structure of the recorded data when assessing the significance of the results. Data was collected in the following nested hierarchies: Neurons, within recording days, within microscope fields of view, within mice. Bootstrapping was performed on each level of this hierarchy 1000 times. The same number of samples as originally recorded are resampled with replacement from the original samples, mimicking a sampling distribution. This resampling is performed at each layer of the hierarchy. For each replicate, summary statistics are calculated on the pooled resampled statistic. Where population level statistics for a session are reported, cells within each session are averaged, and then higher level resampling is performed, as normal. SEM values reported are from the resampled population.

As in Prince et al. (2021), we calculated *p* values as the proportion of pairs of replicates where the statistic for group A exceeded that of group B. This is a direct probability *P* of the population of A being greater than or equal to the population of B and equates to the proportion of the joint probability distribution that falls above the *x* = *y* line. This is in contrast with *p* values used in statistical tests such as the *t*-test or ANOVA, which indicate the probability of obtaining results as extreme as those observed, given that the null hypothesis is true. The probability is significant if *P >* (1 − *α/*2) or if *P < α/*2, where *α* is the significance level. *P >* (1 −*α/*2) denotes that the probability that the WT mean is greater than the 5xFAD mean is significant while *P <* (*α/*2) denotes that the probability that the WT mean is less than the 5xFAD mean is significant. We report bootstrap *p* values simply as *p* for simplicity and for cases where the probability that the WT mean is greater than the 5xFAD mean is significant, we report 1 −*p* to avoid confusion with standard *p* values (Wallace et al., 2020). We report significance values as follows: ^+^*p <* 0.05 (*α* = 0.10), \**p <* 0.025 (*α* = 0.05), \*\**p <* 0.005 (*α* = 0.01), \*\*\**p <* 0.0025 (*α* = 0.005). Throughout the paper, we report mean *±* SEM where the mean is calculated from the actual data and the SEM from the resampled population.

For comparison, we report statistics obtained from Kruskal-Wallis test with post hoc Dunn’s test and effect sizes for all results in Table S1.

#### Estimation statistics and Cumming plots

Estimation statistics reports the magnitude of the effect size and its confidence interval in contrast to significance testing. We used the Cumming plot from the DABEST framework (Ho et al., 2020) to visualize the effect size and to show comparison between multiple groups. The upper part of the plot shows the distribution of raw data with the mean *±* SEM. The lower part shows the median difference, with 95% confidence interval, between a group and the left-most group which is taken as the reference. The median difference is calculated from 5000 bootstrap resamples with replacement.

### Data availability

Data and code to generate the figures from this paper will be made available at DataDryad, with DOI to be inserted here at the time of publication. The raw dataset reported in this paper will be shared by the Lead contact upon reasonable request. Any additonal information required to reanalyze the data reported in this paper is available from the Lead contact upon reasonable request.

### Supplemental Information

Document S1. Figures S1-S8. Table S1.

## Supplemental Information

**Figure S1.**
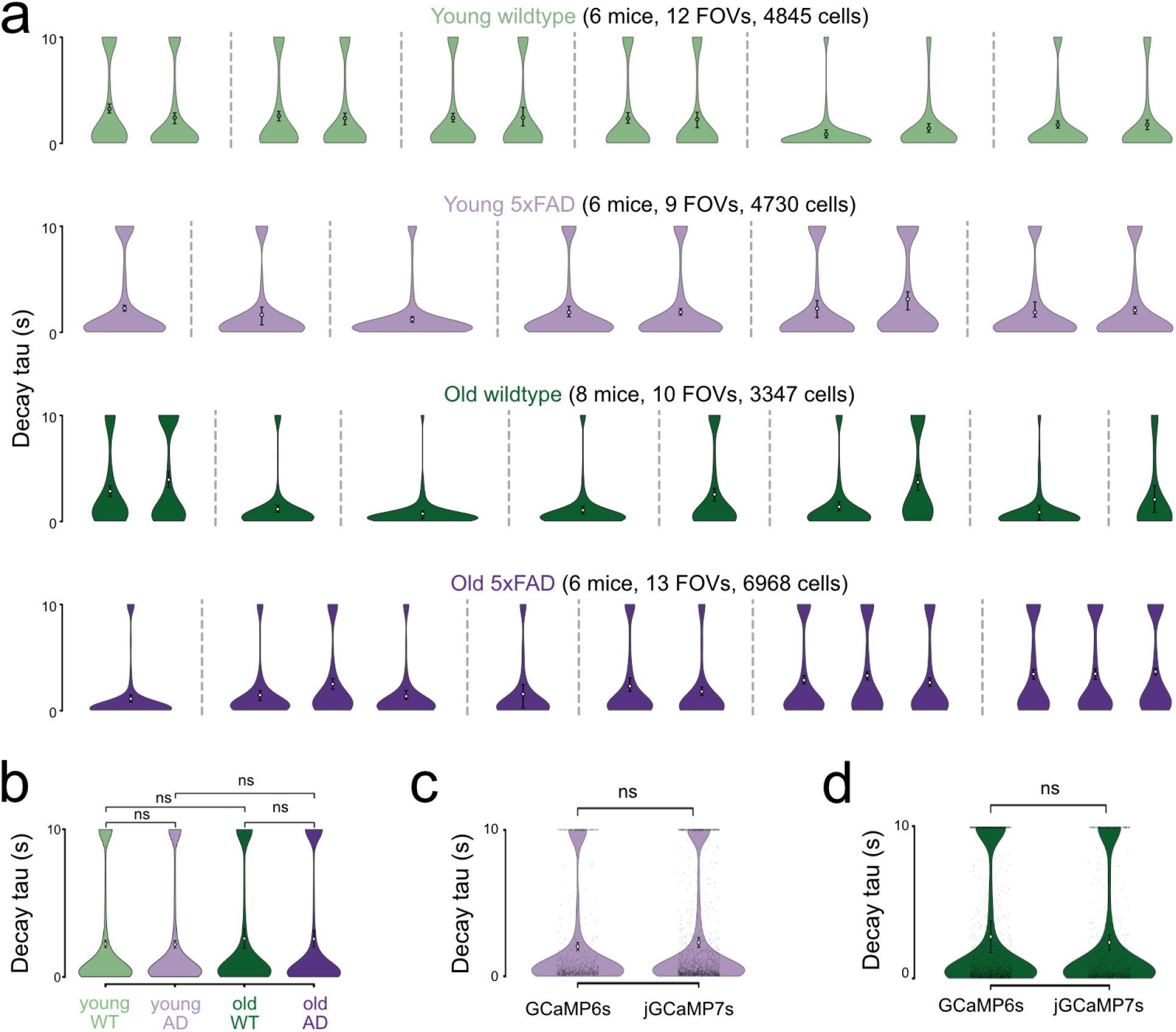
Calcium transient decay constant is comparable across groups and across GCaMP versions. **a** Cell-level decay time constants of CA1 calcium transients are shown for young WT, young 5xFAD, old WT and old 5xFAD mice. Each violin represents one field of view (FOV) from one mouse. For each cell, isolated OASIS events were aligned to their event peaks, df/f traces from 1 s before to 4 s after each event were extracted. Traces were baseline-subtracted using the median pre-event df/f signal, averaged across events and fitted over the post-event period with an exponential model *Ae*^−*t*/*τ*^ + *C*. The fitted *τ* was used as the cell decay time constant. Black points and error bars indicate the FOV mean and 95% bootstrap confidence interval. The number of mice, FOVs and cells are indicated above each group. **b** Group-level comparison of cell decay time constants across young WT, young 5xFAD, old WT and old 5xFAD mice. Statistical comparisons were performed using hierarchical bootstrap analysis; ns: not significant. **c**,**d** Decay time constants compared between GCaMP6s and jGCaMP7s in young 5xFAD mice (**c**) and old WT mice (**d**).

**Figure S2.**
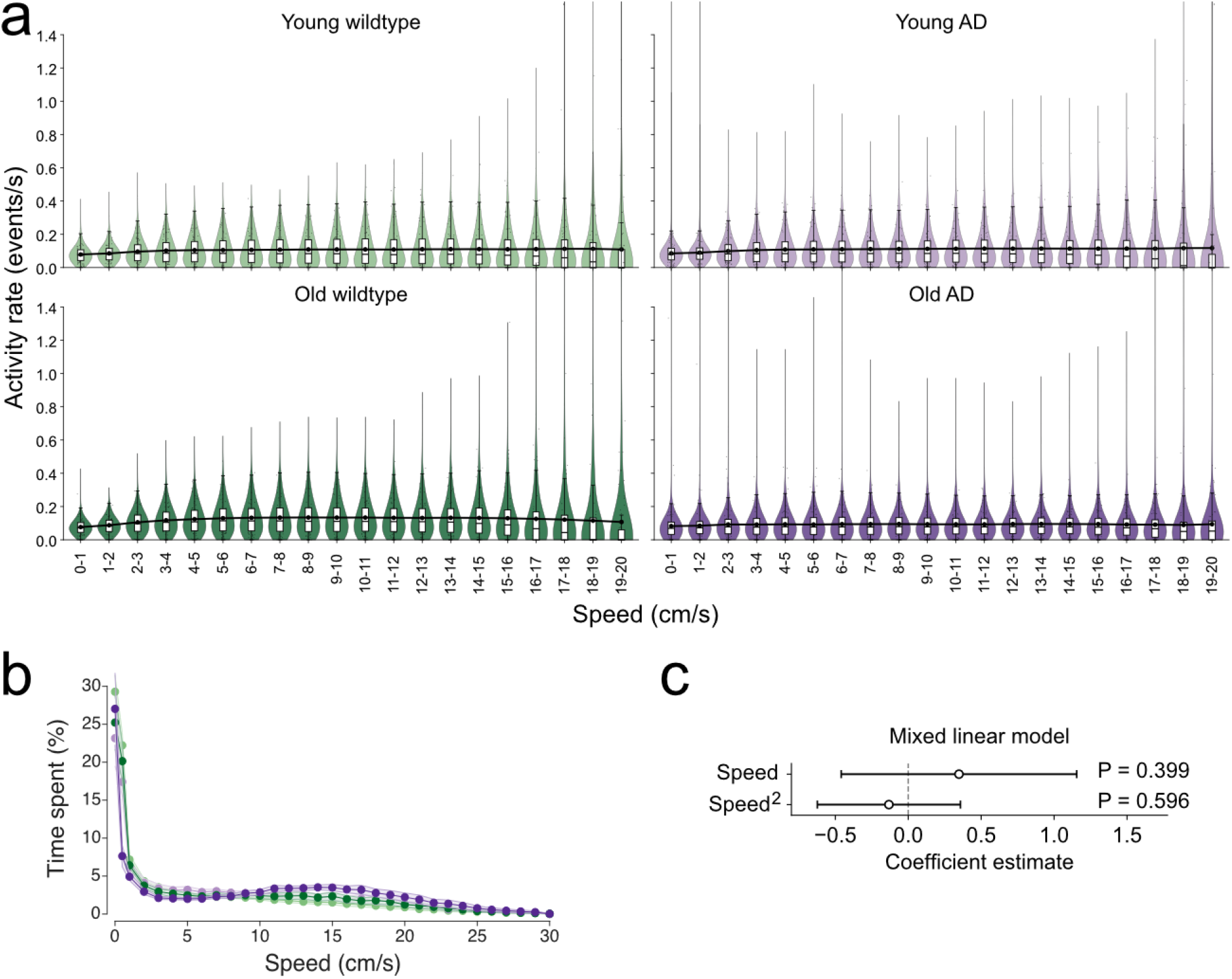
Speed has no significant effect on dynamic range. **a** Activity rate distributions across running speed bins. Event rate of CA1 neurons as a function of running speed, shown separately for young WT (WTY), young 5xFAD (ADY), old WT (WTO), and old 5xFAD (ADO) mice. Violin plots show the distribution of neuronal event rates within each speed bin; overlaid boxplots indicate the median and interquartile range. Black dots and connecting lines show the mean event rate across speed bins for each group. **b** Time spent by animal per speed bin, averaged across all animals in a group. Analysis of dynamic range was limited to 20 cm/s to eliminate speed bins with very low occupancy for some groups. Color codes as in (a). **c** Mixed linear model testing whether speed explains variation in dynamic range. Cell-level dynamic range is modelled as a function of speed and squared speed. Points show coefficient estimates and horizontal bars show 95% confidence intervals; the dashed vertical line marks zero effect. Neither the linear nor quadratic speed term significantly predicted dynamic range, indicating that the observed dynamic range differences were not explained by speed.

**Figure S3:**
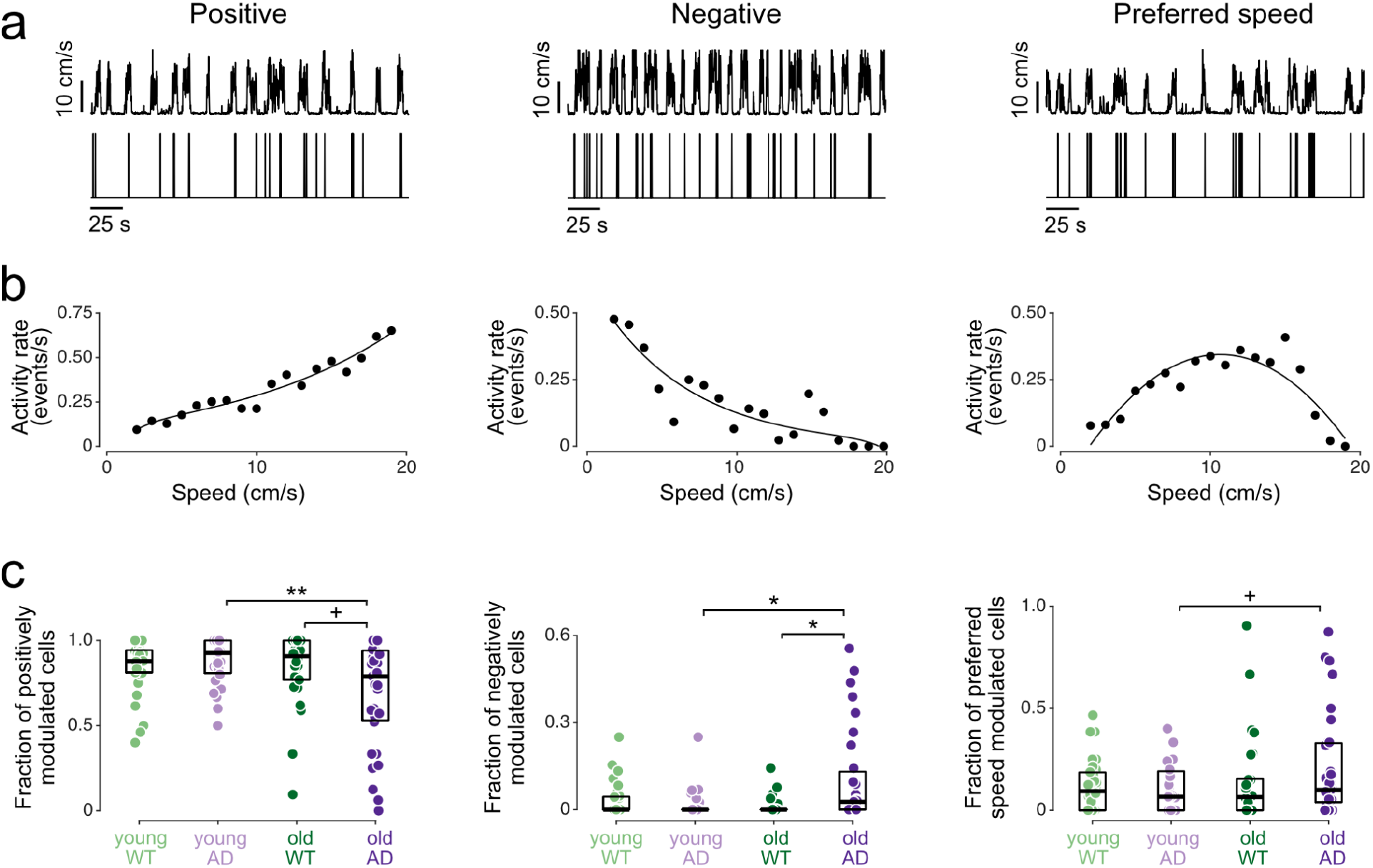
Subtype-specific differences in proportions of speed-modulated cells in 5xFAD mice. **a** Instantaneous speed (top) and firing rate (bottom) for examples of three different types of speed-modulated cells. **b** Speed tuning curves for the different speed-modulated cells in (**a**). Solid lines denote best fit lines. **c** Proportion of speed-modulated cells falling in each of the three classes. We observed significantly less positively-modulated cells and more negatively-modulated cells in old 5xFAD mice compared to old WT and young 5xFAD mice. There were also more preferred-speed cells in old 5xFAd compared to young 5xFAD mice. ^+^*p*<0.05, \**p<*0.025, \*\**p*<0.005. All statistics were performed with hierarchical bootstrap analysis. For all panels, young WT: n = 5,310 cells pooled from 6 animals, 33 sessions; young 5xFAD: n = 4,955 cells, 6 animals, 27 sessions; old WT: n = 3,517 cells, 8 animals, 27 sessions; old 5xFAD: n = 7,400 cells, 6 animals, 31 sessions.

**Figure S4.**
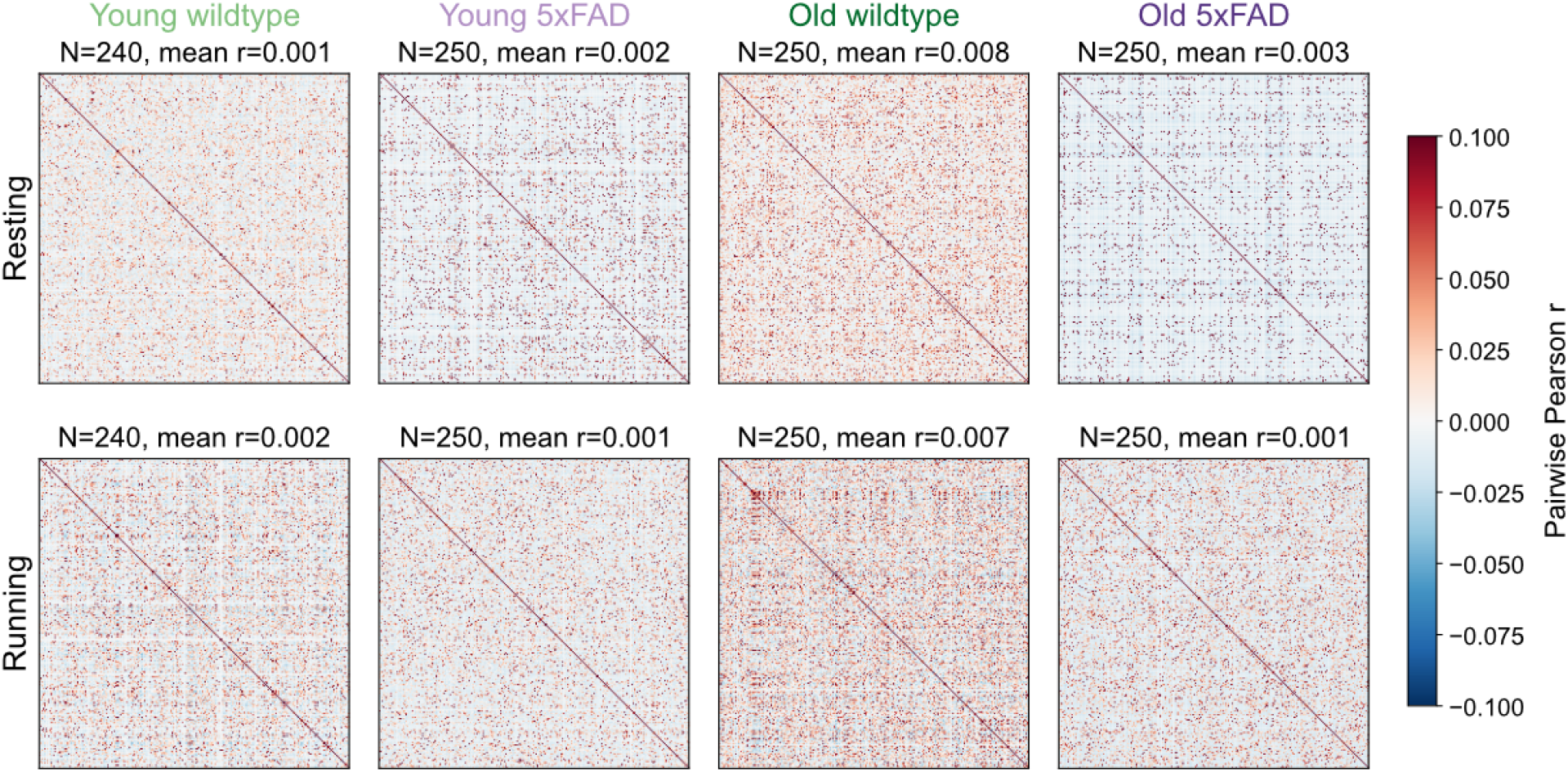
Correlation matrices for all cell pairs in a field of view for a representative mouse from each group. Pairwise Pearson correlation matrices of CA1 neuronal activity are shown for one representative field of view from each group: young WT, young 5xFAD, old WT and old 5xFAD mice. Matrices are shown separately for resting and running periods. For each representative FOV, 250 neurons were randomly selected where available; if fewer than 250 neurons were present, all available neurons were included. Color indicates the pairwise Pearson correlation coefficient. The number of neurons included and the mean pairwise correlation are indicated above each matrix.

**Figure S5.**
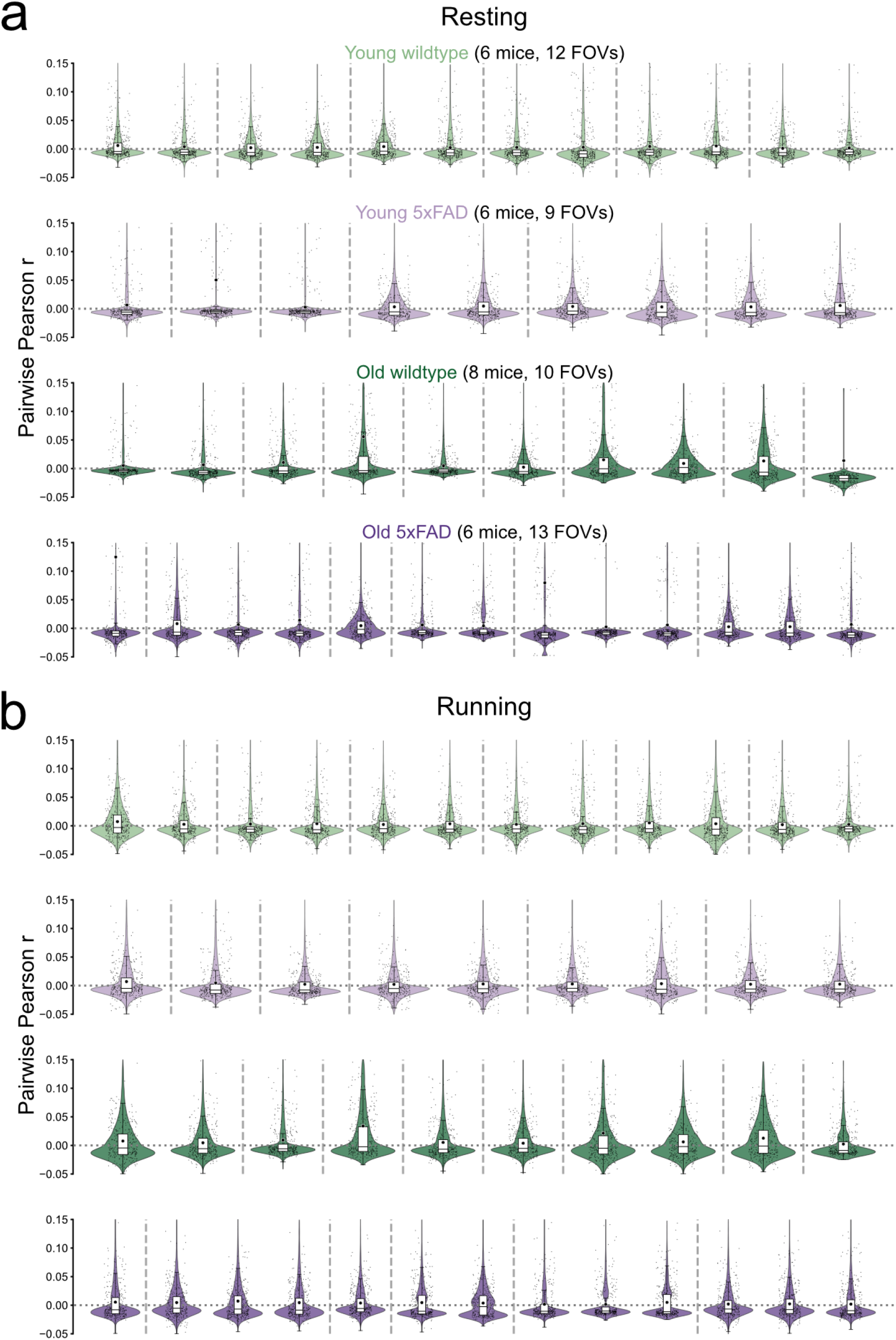
Distributions of pairwise correlations for synchrony calculation. **a** Resting-state pairwise correlation distributions across mice. Pairwise Pearson correlations between CA1 neurons during resting periods are shown separately for young WT, young 5xFAD, old WT and old 5xFAD mice within different field of views (FOVs). Each violin represents the distribution of pairwise correlations within one mouse FOV; sparse black points show sampled neuron pairs.Overlaid boxplots indicate the median and interquartile range, and black dots show the mean pairwise correlation for each FOV. The horizontal dotted line marks zero correlation. The number of mice and FOVs included in each group is indicated above each panel. **b** Running-state pairwise correlation distributions across mice. Same as **a**, but for running periods.

**Figure S6.**
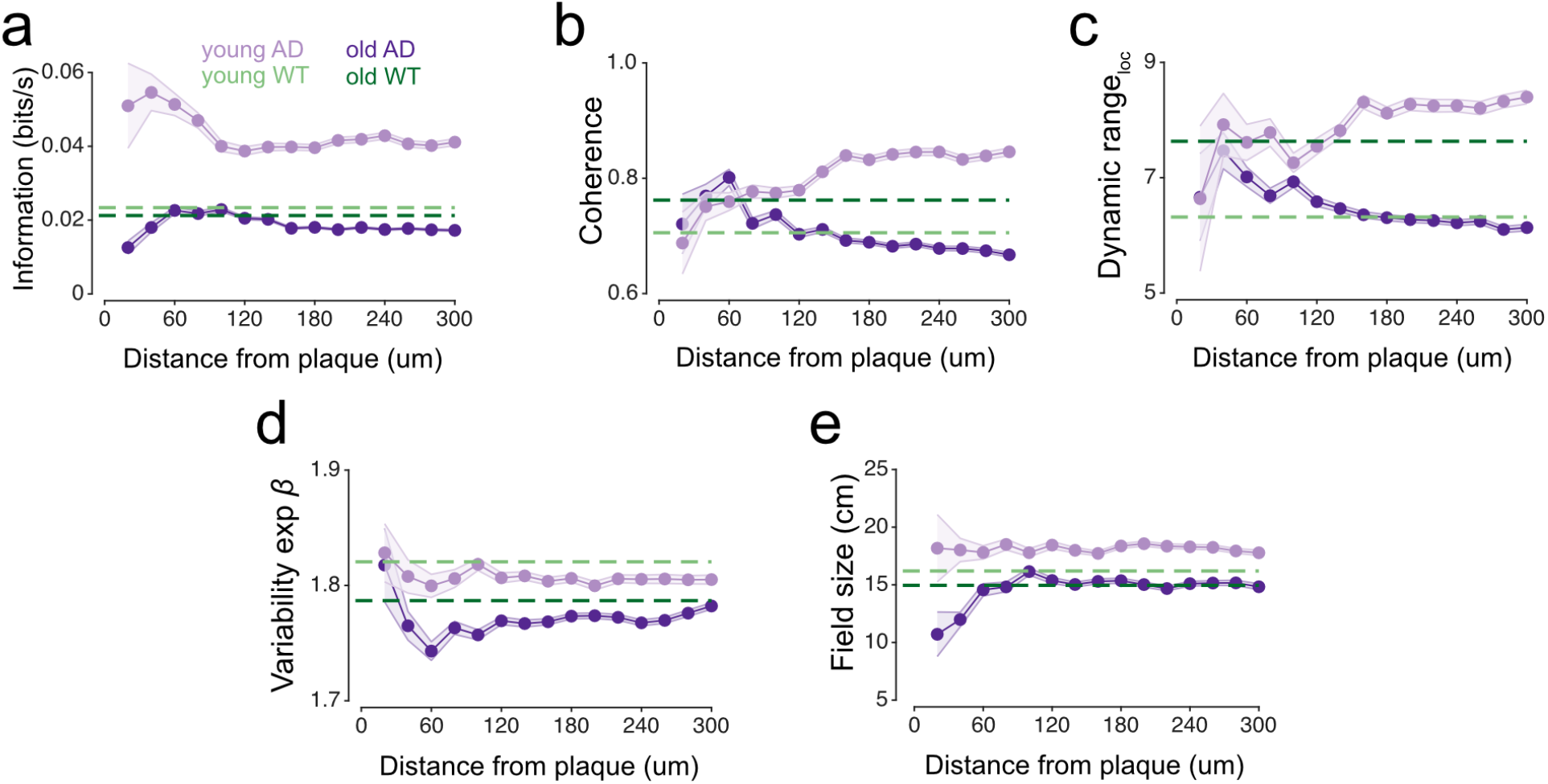
Poorer spatial coding near plaques in old AD mice. Average (**a**) spatial information, (**b**) coherence, (**c**) dynamic range of activity in response to location, (**d**) variability exponent, and (**e**) place field size as a function of distance from plaque. Error bars show SEM. For a-f, young WT: n = 5082 cells pooled from 6 animals, 31 sessions; young 5xFAD: 4955 cells, 6 animals, 27 sessions; old WT: 3268 cells, 8 animals, 24 sessions; old 5xFAD: 7400 cells, 6 animals, 31 sessions.

**Figure S7.**
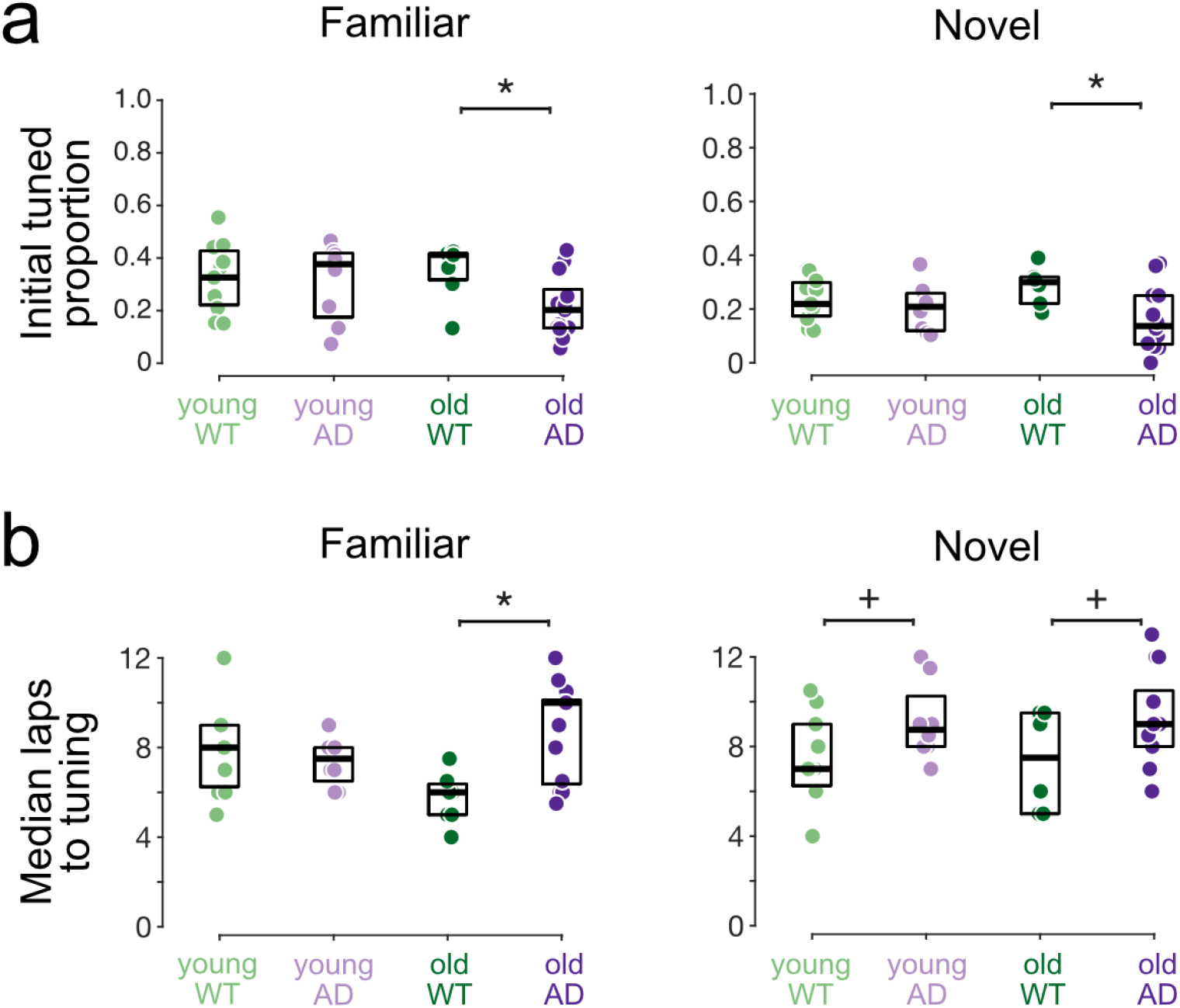
AD mice show slower rate of tuning acquisition in novel environments. **(a)** Proportion of place cells appearing initially. \**p*<0.01. **(b)** Number of laps until tuning for place cells that were not initially tuned. ^+^*p*<0.05, \**p=*0.01. All statistics were performed with hierarchical bootstrap analysis. For all panels, young WT: n = 449 place cells pooled from 6 animals, 11 sessions; young 5xFAD: n = 390 place cells, 5 animals, 8 sessions; old WT: n = 280 place cells, 6 animals, 6 sessions; old 5xFAD: n = 291 place cells, 5 animals, 8 sessions.

**Figure S8.**
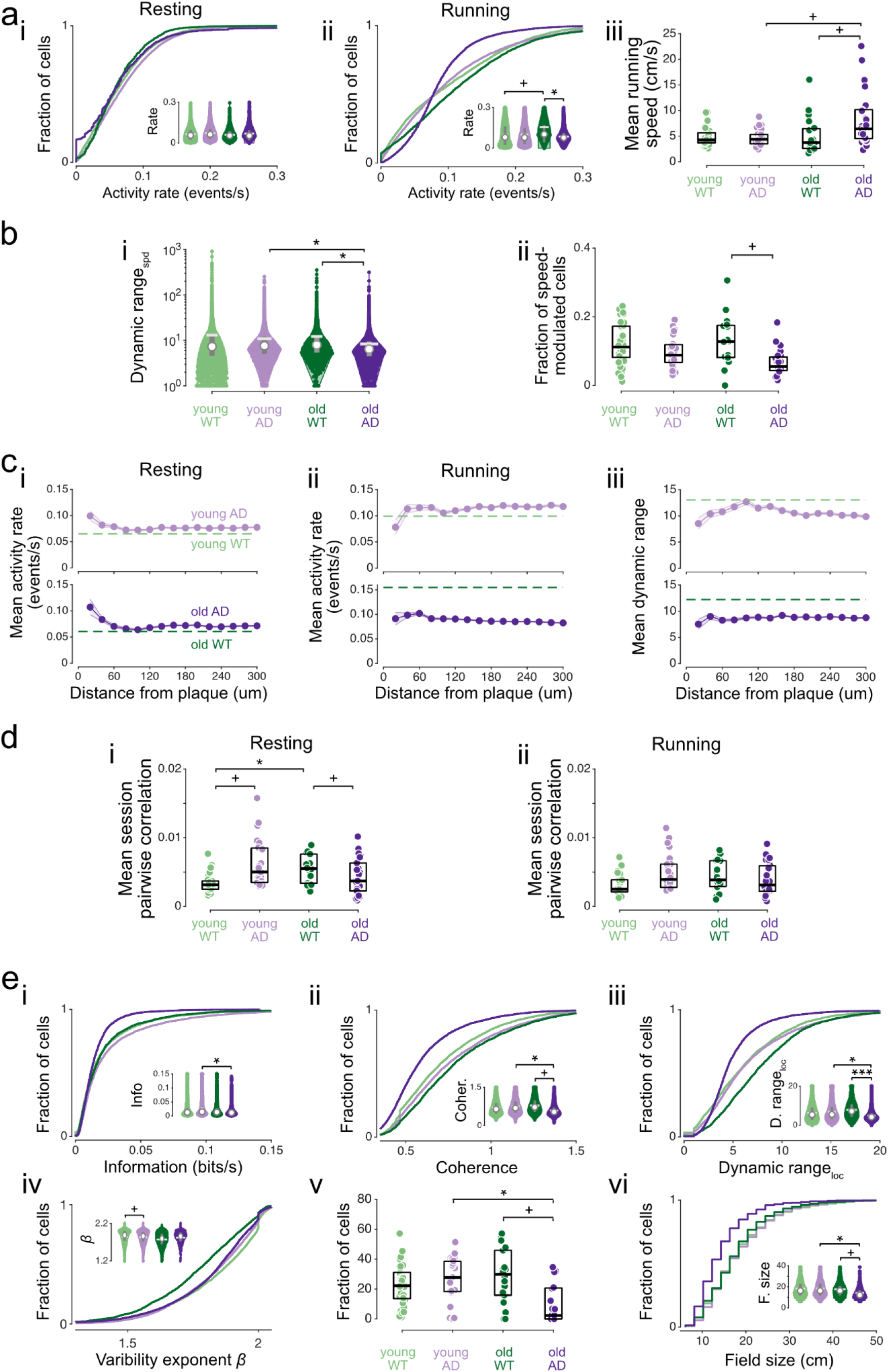
Main results do not change when animals older than 9 months are excluded. **a** Cumulative histograms and violin plots (inset) of activity rate distributions during (**i**) resting and (**ii**) running periods. Color codes as in (**iii**). Violin plots show mean (white solid line), median (white circle) and interquartile range (gray vertical line). **iii** Average running speed during the first 50 laps of each session. Each data point indicates one running session. Boxplot edges denote quartiles while black bars denote median. **b** (**i**) Violin plots of dynamic range of neuronal activity during locomotion. (**ii**) Proportion of cells in each group showing significant speed modulation. Each data point indicates a session. **c** (**i-iii**) Average neuronal activity during (**i**) resting and (**ii**) running periods, and (**iii**) average speed-related dynamic range versus distance from plaques. Dashed lines in (**i-iii**) denote the average for the corresponding age-matched WT group. SEM lines are shown. **d** Average pairwise Pearson correlation for during (**i**) resting and (**ii**) running periods. Each data point indicates an imaging session. **e** Cumulative histograms and violin plots (insets) of (**i**) spatial information, (**ii**) coherence, (**iii**) spatial response dynamic range, (**iv**) variability exponent *β* and y-intercept *α*, (**v**) fraction of place cells and (**vi**) place field size. For **a-e**, young WT: n = 5,168 cells pooled from 6 animals, 33 sessions; young 5xFAD: n = 4,925 cells, 6 animals, 27 sessions; old WT: n = 2,428 cells, 5 animals, 18 sessions; old 5xFAD: n = 6,378 cells, 5 animals, 24 sessions. All statistics were performed with hierarchical bootstrap analysis. ^+^*p* < 0.050, \**p* < 0.025, \*\*\**p* < 0.0025.

**Table S1.** Statistical significance testing using Kruskal-Wallis test with post hoc Dunn’s test. Effect sizes are given for statistically significant pairs (*p*<0.05; small effect: [0.1,0.3], medium effect: [0.3,0.5], large effect: >0.5).

|  | Young WT vs Young AD |  | Old WT vs Old AD |  | Young WT vs Old WT |  | Young AD vs Old AD |  |
| --- | --- | --- | --- | --- | --- | --- | --- | --- |
|  | *p* value | Effect size | *p* value | Effect size | *p* value | Effect size | *p* value | Effect size |
| **Figure 1** |  |  |  |  |  |  |  |  |
| Resting calcium event rate | <0.001 | 0.06 | 0.312 |  | <0.001 | 0.07 | <0.001 | 0.06 |
| Running calcium event rate | 0.999 |  | <0.001 | 0.06 | <0.001 | 0.07 | 1.000 |  |
| Running speed | 1.000 |  | <0.001 | 0.46 | 0.869 |  | 0.038 | 0.27 |
| **Figure 2** |  |  |  |  |  |  |  |  |
| Dynamic range for speed | 0.006 | 0.028 | <0.001 | 0.062 | <0.001 | 0.065 | <0.001 | 0.06 |
| Fraction of speed-modulated cells | 0.637 |  | 0.048 | 0.260 | 1.000 |  | 0.707 |  |
| **Figure 4** |  |  |  |  |  |  |  |  |
| Resting synchrony | 0.034 | 0.27 | 0.580 |  | <0.001 | 0.56 | 0.989 |  |
| Running synchrony | 0.998 |  | 0.004 | 0.37 | 0.001 | 0.43 | 0.962 |  |
| **Figure 5** |  |  |  |  |  |  |  |  |
| Spatial information | <0.001 | 0.08 | 0.012 | 0.02 | 0.485 |  | <0.001 | 0.06 |
| Coherence | <0.001 | 0.08 | <0.001 | 0.06 | <0.001 | 0.07 | <0.001 | 0.06 |
| Dynamic range for location | <0.001 | 0.05 | <0.001 | 0.06 | <0.001 | 0.07 | <0.001 | 0.06 |
| Variability exponent | <0.001 | 0.05 | <0.001 | 0.05 | <0.001 | 0.07 | <0.001 | 0.05 |
| Fraction of place cells | 0.903 |  | 0.062 | 0.25 | 1.000 |  | 0.003 | 0.39 |
| Field size | <0.001 | 0.04 | <0.001 | 0.06 | <0.001 | 0.07 | <0.001 | 0.06 |
| **Figure 6** |  |  |  |  |  |  |  |  |
| Median laps to tuning FAM | 0.958 |  | <0.001 | 0.14 |  |  |  |  |
| Median laps to tuning NOV | 0.005 | 0.09 | <0.001 | 0.10 |  |  |  |  |

## References

Aghajan, Z.M., Acharya, L., Moore, J., Cushman, J., Vuong, C., Mehta, M.R., 2015. Impaired spatial selectivity and intact phase precession in two-dimensional virtual reality. Nat Neurosci 18, 121–128.

Bezzina, C., Verret, L., Juan, C., Remaud, J., Halley, H., Rampon, C., Dahan, L., 2015. Early onset of hypersynchronous network activity and expression of a marker of chronic seizures in the tg2576 mouse model of alzheimer’s disease. PLoS ONE 10, e0119910.

Broussard, J.I., Redell, J.B., Maynard, M.E., Zhao, J., Moore, A., Mills, R.W., Hood, K.N., Underwood, E., Roysam, B., Dash, P.K., 2022. Impaired experience-dependent refinement of place cells in a rat model of alzheimer’s disease. J Alzheimers Dis 86, 1907–1916.

Busche, M.A., Chen, X., Henning, H.A., Reichwald, J., Staufenbiel, M., Sakmann, B., Konnerth, A., 2012. Critical role of soluble amyloid-β for early hippocampal hyperactivity in a mouse model of alzheimer’s disease. Proceedings of the National Academy of Sciences 109, 8740–8745.

Busche, M.A., Eichhoff, G., Adelsberger, H., Abramowski, D., Wiederhold, K.H., Haass, C., Staufenbiel, M., Konnerth, A., Garaschuk, O., 2008. Clusters of hyperactive neurons near amyloid plaques in a mouse model of alzheimer’s disease. Science 321, 1686–1689.

Cacucci, F., Yi, M., Wills, T.J., Chapman, P., O’Keefe, J., 2008. Place cell firing correlates with memory deficits and amyloid plaque burden in tg2576 alzheimer mouse model. Proceedings of the National Academy of Sciences 105, 7863–7868.

Carandini, M., 2004. Amplification of trial-to-trial response variability by neurons in visual cortex. PLoS biology 2, e264.

Cheng, J., Ji, D., 2013. Rigid firing sequences undermine spatial memory codes in a neurodegenerative mouse model. eLife 2, e00647.

Chockanathan, U., Padmanabhan, K., 2024. Differential disruptions in population coding along the dorsal-ventral axis of ca1 in the app/ps1 mouse model of aβ pathology. PLoS Comput Biol 20, e1012085.

Coughlan, G., Laczó, J., Hort, J., Minihane, A.M., Hornberger, M., 2018. Spatial navigation deficits—overlooked cognitive marker for preclinical alzheimer disease? Nature Reviews Neurology 14, 496–506.

Fortel, I., Zhan, L., Ajilore, O., Wu, Y., Mackin, S., Leow, A., 2023. Disrupted excitation-inhibition balance in cognitively normal individuals at risk of alzheimer’s disease. Journal of Alzheimer’s Disease 95, 1449–1467.

Friedrich, J., Yang, W., Soudry, D., Mu, Y., Ahrens, M.B., Yuste, R., Peterka, D.S., Paninski, L., 2017. Multi-scale approaches for highspeed imaging and analysis of large neural populations. PLOS Comput. Biol. 13, e1005685. doi:10.1371/journal.pcbi.1005685.

Giovannucci, A., Friedrich, J., Gunn, P., Kalfon, J., Brown, B.L., Koay, S.A., Taxidis, J., Najafi, F., Gauthier, J.L., Zhou, P., Khakh, B.S., Tank, D.W., Chklovskii, D.B., Pnevmatikakis, E.A., 2019. CaImAn an open source tool for scalable calcium imaging data analysis. eLife 8, 38173. doi:10.7554/eLife.38173.

Go, M.A., Rogers, J., Gava, G.P., Davey, C.E., Prado, S., Liu, Y., Schultz, S.R., 2021. Place cells in head-fixed mice navigating a floating real-world environment. Frontiers in Cellular Neuro-science 15, 19.

Góis, Z.H.T.D., Tort, A.B.L., 2018. Characterizing speed cells in the rat hippocampus. Cell Reports 25, 1872–1884.

Gollo, L.L., Kinouchi, O., Copelli, M., 2009. Active dendrites enhance neuronal dynamic range. PLoS Computat Biol 5, e1000402.

Gollo, L.L., Mirasso, C., Eguíluz, V.M., 2012. Signal integration enhances the dynamic range in neuronal systems. Phys Rev E 85, 040902.

Grutzendler, J., Helmin, K., Tsai, J., Gan, W., 2007. Various dendritic abnormalities are associated with fibrillar amyloid deposits in alzheimer’s disease. Ann N Y Acad Sci 1097, 30–39.

Hainmueller, T., Bartos, M., 2018. Parallel emergence of stable and dynamic memory engrams in the hippocampus. Nature 558, 292–296.

Haj, M.E., Robin, F., 2021. Repeated recall on source misattribution in alzheimer’s disease. Memory 29, 1354–1361.

Harvey, C.D., Collman, F., Dombeck, D.A., Tank, D.W., 2009. Intra-cellular dynamics of hippocampal place cells during virtual navigation. Nature 461, 941–946.

Ho, J., Tumkaya, T., Aryal, S., Choi, H., Claridge-Chang, A., 2020. Moving beyond p values: data analysis with estimation graphics. Nature Methods 16, 565–566.

Hyman, B.T., Van Hoesen, G.W., Damasio, A.R., Barnes, C.L., 1984. Alzheimer’s disease: cell-specific pathology isolates the hippocampal formation. Science 225, 1168–1170.

Iaccarino, H.F., Singer, A.C., Martorell, A.J., Rudenko, A., Gao, F., Gillingham, T.Z., Mathys, H., Seo, J., Kritskiy, O., Abdurrob, F., Adaikkan, C., Canter, R.G., Rueda, R., Brown, E.N., Boyden, E.S., Tsai, L.H., 2016. Gamma frequency entrainment attenuates amyloid load and modifies microglia. Nature 540, 230–235.

Jawhar, S., Trawicka, A., Jenneckens, C., Bayer, T.A., Wirths, O., 2012. Motor deficits, neuron loss, and reduced anxiety coinciding with axonal degeneration and intraneuronal aβ aggregation in the 5xfad mouse model of alzheimer’s disease. Neurobiology of aging 33, 196–e29.

Jun, H., Bramian, A., Soma, S., Saito, T., Saido, T.C., Igarashi, K.M., 2020. Disrupted place cell remapping and impaired grid cells in a knockin model of alzheimer’s disease. Neuron 107, 1095–1112.

Kropff, E., Carmichael, J.E., Moser, M.B., Moser, E.I., 2015. Speed cells in the medial entorhinal cortex. Nature 523, 419–424. doi:10.1038/nature14622.

Lagadeca, S., Rotureaua, L., Hémarb, A., Macreza, N., Delcassoa, S., Jeanteta, Y., Cho, Y.H., 2012. Early temporal short-term memory deficits in double transgenic app/ps1 mice. Neurobiol Aging 33, 203.e1–203.e11.

Langston, R.F., Ainge, J.A., Couey, J.J., Canto, C.B., Bjerknes, T.L., Witter, M.P., Moser, E.I., Moser, M.B., 2010. Development of the spatial representation system in the rat. Science 328, 1576–1580.

Le, R., Cruz, L., Urbanc, B., Knowles, R.B., Hsiao-Ashe, K., Duff, K., Irizzary, M.C., Stanley, H.E., Hyman, B.T., 2001. Plaqueinduced abnormalities in neurite geometry in transgenic models of alzheimer disease: Implications for neural system disruption. J Neuropathol Exp Neurol 60, 735–758.

Lin, X., Chen, L., Baglietto-Vargas, D., Kamalipour, P., Ye, Q., LaFerla, F.M., Nitz, D.A., Holmes, T.C., Xu, X., 2022. Spatial coding defects of hippocampal neural ensemble calcium activities in the triple-transgenic alzheimer’s disease mouse model. Neurobiol Disease 162, 105562.

Mably, A.J., Gereke, B.J., Jones, D.T., Colgin, L.L., 2017. Impairments in spatial representations and rhythmic coordination of place cells in the 3xtg mouse model of alzheimer’s disease. Hippocampus 27, 378–392.

Moolman, D.L., Vitolo, O.V., Vonsattel, J.P.G., Shelanski, M.L., 2004. Dendrite and dendritic spine alterations in alzheimer models. J Neurocytol 33, 377–387.

Muller, R.U., Kubie, J.L., 1987. The effects of changes in the environment on the spatial firing of hippocampal complex-spike cells. Journal of Neuroscience 7, 1951–1968.

Muzzu, T., Mitolo, S., Gava, G.P., Schultz, S.R., 2018. Encoding of locomotion kinematics in the mouse cerebellum. PLOS ONE 13, e0203900.

Oakley, H., Cole, S.L., Logan, S., Maus, E., Shao, P., Craft, J., Guillozet-Bongaarts, A., Ohno, M., Disterhoft, J., Van Eldik, L., et al., 2006. Intraneuronal β-amyloid aggregates, neurodegeneration, and neuron loss in transgenic mice with five familial alzheimer’s disease mutations: potential factors in amyloid plaque formation. Journal of Neuroscience 26, 10129–10140.

O’Keefe, J., Dostrovsky, J., 1971. The hippocampus as a spatial map: Preliminary evidence from unit activity in the freely-moving rat. Brain Research 34, 171–175.

Ólafsdóttir, H.F., Bush, D., Barry, C., 2018. The role of hippocampal replay in memory and planning. Current Biology 28, R37–R50.

Palop, J.J., Chin, J., Roberson, E.D., Wang, J., Thwin, M.T., Bien-Ly, N., Yoo, J., Ho, K.O., Yu, G.Q., Kreitzer, A., et al., 2007. Aberrant excitatory neuronal activity and compensatory remodeling of inhibitory hippocampal circuits in mouse models of alzheimer’s disease. Neuron 55, 697–711.

Palop, J.J., Mucke, L., 2016. Network abnormalities and interneuron dysfunction in alzheimer disease. Nature Reviews Neuroscience 17, 777–792.

Poll, S., Mittag, M., Musacchio, F., Justus, L.C., Giovannetti, E.A., Steffen, J., Wagner, J., Zohren, L., Schoch, S., Schmidt, B., Jack-son, W.S., Ehninger, D., Fuhrmann, M., 2020. Memory trace interference impairs recall in a mouse model of alzheimer’s disease. Nat Neurosci 23, 952–958.

Prince, S.M., Paulson, A.L., Jeong, N., Zhang, L., Amigues, S., Singer, A.C., 2021. Alzheimer’s pathology causes impaired inhibitory connections and reactivation of spatial codes during spatial navigation. Cell Reports 35, 109008.

Saravanan, V., Berman, G.J., Sober, S.J., 2020. Application of the hierarchical bootstrap to multi-level data in neuroscience. Neuron Behav Data Anal Theory 3.

Selkoe, D.J., 2001. Alzheimer’s disease: genes, proteins, and therapy. Physiological Reviews 81, 741–766.

Shah, D., Praet, J., Hernandez, A.L., Höfling, C., Anckaerts, C., Bard, F., Morawski, M., Detrez, J.R., Prinsen, E., Villa, A., Vos, W.H.D., Maggi, A., D’Hooge, R., Balschun, D., Rossner, S., Verhoye, M., der Linden, A.V., 2016. Early pathologic amyloid induces hypersynchrony of bold resting-state networks in transgenic mice and provides an early therapeutic window before amyloid plaque deposition. Alzheimer’s & Dementia 12, 964–976.

Skaggs, W., Mcnaughton, B., Gothard, K., 1992. An information-theoretic approach to deciphering the hippocampal code. Advances in neural information processing systems 5.

Skaggs, W.E., McNaughton, B.L., Wilson, M.A., Barnes, C.A., 1996. Theta phase precession in hippocampal neuronal populations and the compression of temporal sequences. Hippocampus 6, 149–172.

Squire, L.R., 2004. Memory systems of the brain: a brief history and current perspective. Neurobiology of Learning and Memory 82, 171–177.

Takamura, R., Mizuta, K., Sekine, Y., Islam, T., Saito, T., Sato, M., Ohkura, M., Nakai, J., Ohshima, T., Saido, T., Hayashi, Y., 2021. Modality-specific impairment of hippocampal ca1 neurons of alzheimer’s disease model mice. Journal of Neuroscience 41, 5315–4329.

Teri, L., Larson, E.B., Reifler, B.V., 1988. Behavioral disturbance in dementia of the alzheimer’s type. Journal of the American Geriatrics Society 36, 1–6.

Tolhurst, D.J., Movshon, J.A., Dean, A.F., 1983. The statistical reliability of signals in single neurons in cat and monkey visual cortex. Vision research 23, 775–785.

Tsai, J., Grutzendler, J., Duff, K., Gan, W., 2004. Fibrillar amyloid deposition leads to local synaptic abnormalities and breakage of neuronal branches. Nat Neurosci 7, 1181–1183.

Šišková, Z., Justus, D., Kaneko, H., Friedrichs, D., Henneberg, N., Beutel, T., Pitsch, J., Schoch, S., Becker, A., von der Kammer, H., Remy, S., 2014. Dendritic structural degeneration is functionally linked to cellular hyperexcitability in a mouse model of alzheimer’s disease. Neuron 84, 1023–1033.

Van Nifterick, A.M., Mulder, D., Duineveld, D.J., Diachenko, M., Scheltens, P., Stam, C.J., Van Kesteren, R.E., Linkenkaer-Hansen, K., Hillebrand, A., Gouw, A.A., 2023. Resting-state oscillations reveal disturbed excitation–inhibition ratio in alzheimer’s disease patients. Scientific reports 13, 7419.

Verret, L., Mann, E.O., Hang, G.B., Barth, A.M., Cobos, I., Ho, K., Devidze, N., Masliah, E., Kreitzer, A.C., Mody, I., Mucke,L., Palop, J.J., 2012. Inhibitory interneuron deficit links altered network activity and cognitive dysfunction in alzheimer model. Cell 149, 708–721.

Wallace, J., Lord, J., Dissing-Olesen, L., Stevens, B., Murthy, V.N., 2020. Microglial depletion disrupts normal functional development of adult-born neurons in the olfactory bulb. eLife 9, e50531.

Wang, L., Wang, Y., long Fu, W., hong Cao, L., 2017. Modulation of neuronal dynamic range using two different adaptation mechanisms. Neural Regen Res 12, 447–451.

Zhang, H., Chen, L., Johnston, K.G., Crapser, J., Green, K.N., Ha, N.M.L., Tenner, A.J., Holmes, T.C., Nitz, D.A., Xu, X., 2023. Degenerate mapping of environmental location presages deficits in object-location encoding and memory in the 5xfad mouse model for alzheimer’s disease. Neurobiology of disease 176, 105939.

Zhang, S., Schönfeld, F., Wiskott, L., Manahan-Vaughan, D., 2014. Spatial representations of place cells in darkness are supported by path integration and border information. Front Behav Neurosci 8, 222.

Zhao, R., Fowler, S.W., Chiang, A.C., Ji, D.,, Jankowsky, J.L., 2014. Impairments in experience-dependent scaling and stability of hippocampal place fields limit spatial learning in a mouse model of alzheimer’s disease. Hippocampus 24, 963–978.

Zong, W., Obenhaus, H.A., Skytøen, E.R., Eneqvist, H., de Jong, N.L., Vale, R., Jorge, M.R., Moser, M.B., Moser, E.I., 2022. Large-scale two-photon calcium imaging in freely moving mice. Cell 185, 1240–1256.

